# Prokaryote communities associated with different types of tissue formed and substrates inhabited by *Serpula lacrymans*

**DOI:** 10.1101/2022.12.23.521786

**Authors:** Julia Embacher, Susanne Zeilinger, Martin Kirchmair, Sigrid Neuhauser

**Author notes:** Corresponding author* Correspondence to Sigrid Neuhauser, Institute of Microbiology, Technikerstrasse 25, A-6020 Innsbruck, Mail. E-mail addresses Julia Embacher, Susanne Zeilinger, Martin Kirchmair, and Sigrid Neuhauser. ***Data availability statement.*** The sequences obtained from each sample were submitted to Sequence Read Archive and are available under BioProject PRJNA878418 (accession numbers SAMN30726529 to SAMN30726547). ***Funding statement.*** This work was funded by Universität Innsbruck (Doctoral program BioApp and ‘Early Stage Funding’ 2021-Bio-1, no. 346314). SN was funded by the Austrian Science Fund: grant Y0801-B16. ***CRediT (Contribution Roles Taxonomy):*** JE: Conceptualization, Resources, Funding acquisition, Investigation (sample preparation and DNA extraction, Amplicon Results analysis), Methodology, Data curation, Visualization, Formal Analysis, Writing – Original Draft preparation, Writing – review & editing, SZ: Conceptualization, Funding acquisition, Supervision, Writing – review & editing, MK: Supervision, Conceptualization, Resources, Writing – review & editing, SN: Supervision, Conceptualization, Investigation (Amplicon Results analysis), Methodology, Validation, Writing – review & editing.

## Abstract

The basidiomycete *Serpula lacrymans* is responsible for timber destruction in houses. Basidiomycetes are known to harbor a diverse but poorly understood microbial community of bacteria, archaea, yeasts, and filamentous fungi in their fruiting bodies. In this study, we used amplicon-sequencing to analyze the abundance and composition of prokaryotic communities associated with fruiting bodies of *S. lacrymans* and compared them to communities of surrounding material to access the ‘background’ community structure. Our findings indicate that bacterial genera cluster depended on sample type, and that the main driver for microbial diversity is specimen, followed by sample origin. The most abundant bacterial phylum identified in the fruiting bodies was Pseudomonadota, followed by Actinomycetota and Bacteroidota. The prokaryote community of the mycelium was dominated by Actinomycetota, Halobacterota, and Pseudomonadota. Actinomycetota was the most abundant phylum in both environment samples (infested timber and underground scree), followed by Bacillota in wood and Pseudomonadota in underground scree. *Nocardioides, Pseudomonas, Pseudonochardia, Streptomyces* and *Rubrobacter* spp. were among others found to comprise the core microbiome of *S. lacrymans* basidiocarps. This research contributes to the understanding of the holobiont *S. lacrymans* and gives hints to potential bacterial phyla important for its development and life style.

**Highlights:**

- The prokaryote communities associated with *S. lacrymans* mycelia and fruiting bodies as well as wood and non-woody substrate form distinct clusters.
- Across all samples 30% of OTU’s were shared (core microbiome) while the number of unique OTUs was small.
- Fruiting bodies (n= 8) of *S. lacrymans* shared a core set of 365 OTU’s, dominated by Actinobacteriodota (44%), Pseudomonadota (28%), and Acidobacteriodota (9%).
- Tissue/sample type is the main factor influencing diversity, followed by sample origin.

## Introduction

Structure, biodiversity and function of prokaryotic communities associated with fungi are increasingly studied (Benucci and Bonito 2016, Bahram, Vanderpool et al. 2018, Liu, Sun et al. 2018, Tarkka, Drigo et al. 2018, Cullings, Stott et al. 2020). Inter-kingdom interactions between fungi and prokaryotes co-occurring in wood cover the full range of symbiotic interactions from mutualism to antagonistic behavior (Greaves 1971, Clausen 1996, Tornberg, Bååth et al. 2003, Boer, Folman et al. 2005, Boddy, Frankland et al. 2007, Yurkov, Krüger et al. 2012, Johnston, Boddy et al. 2016). Microorganisms that co-exist in the same environment affect each other (e.g. metabolically or physiologically). Examples for specialized interactions are bacteria which can promote hyphal growth (Li, Chen et al. 2016, Schulz-Bohm, Tyc et al. 2017, Oh, Kim et al. 2018), or bacteria which induce fungal metabolite production (Vahdatzadeh, Deveau et al. 2015, Tauber, Gallegos-Monterrosa et al. 2018). However, there is also competition between prokaryotes and fungi for nutrients (Folman, Gunnewiek et al. 2008) and even mycophagy may occur, where prokaryotes feed on living (mycoparasitism) or dead (saprotrophy) fungal material.

Wood destroying fungi are responsible for substantial economic damage, especially in buildings. Their ecology and interaction with other microorganisms in the indoor environment, however, is so far not well understood. Because they destabilize wooden structures of buildings, fungi like *Coniophora puteana, Antrodia* spp., and *Serpula lacrymans* need to be removed from construction timber or - in case of *S. lacrymans* - also from the surrounding masonry via costly restoration procedures [DIN 68800-4 (2020) or Ö-Norm B3802-4 (2015)]. The dry rot fungus *S. lacrymans* is specialized on the degradation of coniferous timber and is characterized by rapid spread and wood degradation. The latter is predominantly affected by temperature and moisture as well as the durability of the wood material (Kržišnik, Lesar et al. 2018). Because of the economic importance (Hyde, Al-Hatmi et al. 2018) and the high effort that is needed to treat buildings in which *S. lacrymans* has been found, its interaction with prokaryotes has been explored to better understand the dynamics of dry rot (Embacher, Neuhauser et al. 2021). Wood colonization by *S. lacrymans* can be supported by bacteria that are modifying wood structure and permeability using biopolymer degrading enzymes (e.g. cellulases and pectinases) that pre-digest the cellulose-microfibrils in timber (Boutelje and Bravery 1968, Rehbein, Koch et al. 2013). Wood has a high carbon content but it is relatively poor in biologically available nitrogen; therefore, an increase in nitrogen caused by nitrogen-fixing bacteria in exchange for carbon has been discussed as beneficial interaction between bacteria and fungi (Cowling and Merrill 2011, Purahong, Wubet et al. 2016, Tláskal, Brabcová et al. 2021). A relatively neutral and unspecific type of interaction is represented by Acidobacteriota, Bacteroidota, and Actinomycetota among others that act as decomposers and that follow on the wood substrate to use the remaining cell wall components once *S. lacrymans* has degraded the cellulose (Berlemont and Martiny 2013, Tláskal and Baldrian 2021, Embacher, Zeilinger et al. in press). Some Pseudomonadota and other prokaryotes are able to obtain nutrients via mycophagy, which from the fungal viewpoint is the negative end of interactions (Tláskal and Baldrian 2021). Bacteria belonging to the genera *Bacillus, Pseudomonas, Alcaligenes, Serratia, Streptomyces, Erwinia*, and *Enterobacteria*, among others, have been shown to play a role in the deterioration of building timber (Boutelje and Bravery 1968, Rogers and Baecker 1991, Dutkiewicz, Sorenson et al. 1992, Gaylarde and Morton 1999, Line 2003, Wójcik-Fatla, Mackiewicz et al. 2022).

Aim of this study was to assess the biodiversity of prokaryotes associated to *S. lacrymans* with culture independent methods. We used amplicon sequencing to study the bacterial community composition in fruiting bodies and mycelial mats of *S. lacrymans* as well as wood and other substrates associated with the dry rot fungus. Differences in the prokaryotic biodiversity between and within samples were analyzed. We aimed to test if (i) the structure of the fruiting bodies’ bacterial communities is dissimilar to that of the surrounding wood- or mineral substrate, but more similar to that of the mycelium; and if (ii) there are any differences in community composition compared to our previous results from a culture-based approach. The obtained results revealed a relatively small group of core OTUs associated with *S. lacrymans* fruiting bodies but characteristic prokaryote taxa associated with each fungal tissue type.

## Material and Methods

### Sample collection and processing

Eight fruiting bodies, two mycelium mats, nine substrate samples (five wood, and four mineral construction material like plaster, brick walls, concrete) were collected from five infested houses in Austria (Tab. 1, Fig. 1, Fig. S1). It was not possible to sample mycelium mats at all sampling points; nonetheless to complete the dataset we included the information from two mycelia mats.

**Tab. 1:** Sampling Information.

| sampling point | Class |  |  |  | Sequence<br>Read Archive<br>no. |
| --- | --- | --- | --- | --- | --- |
|  | fruiting body | mycelium<br>mats | wooden<br>substrate | mineral<br>substrate |  |
| Zirl (SZ) | 1 |  | 1 |  | SAMN30726529,<br>SAMN30726530 |
| Reith im<br>Alpbachtal<br>(SM) | 2 |  | 1 | 1 | SAMN30726531,<br>SAMN30726532,<br>SAMN30726533,<br>SAMN30726543 |
| Wolfsegg am<br>Hausruck<br>(WE) | 2 |  | 1 | 1 | SAMN30726534,<br>SAMN30726535,<br>SAMN30726536,<br>SAMN30726537, |
| Grins (G) | 2 | 1 | 1 | 1 | SAMN30726538,<br>SAMN30726539,<br>SAMN30726540,<br>SAMN30726541,<br>SAMN30726542, |
| Tarrenz (T) | 1 | 1 | 1 | 1 | SAMN30726544,<br>SAMN30726545,<br>SAMN30726546,<br>SAMN30726547 |

**Figure 1:**
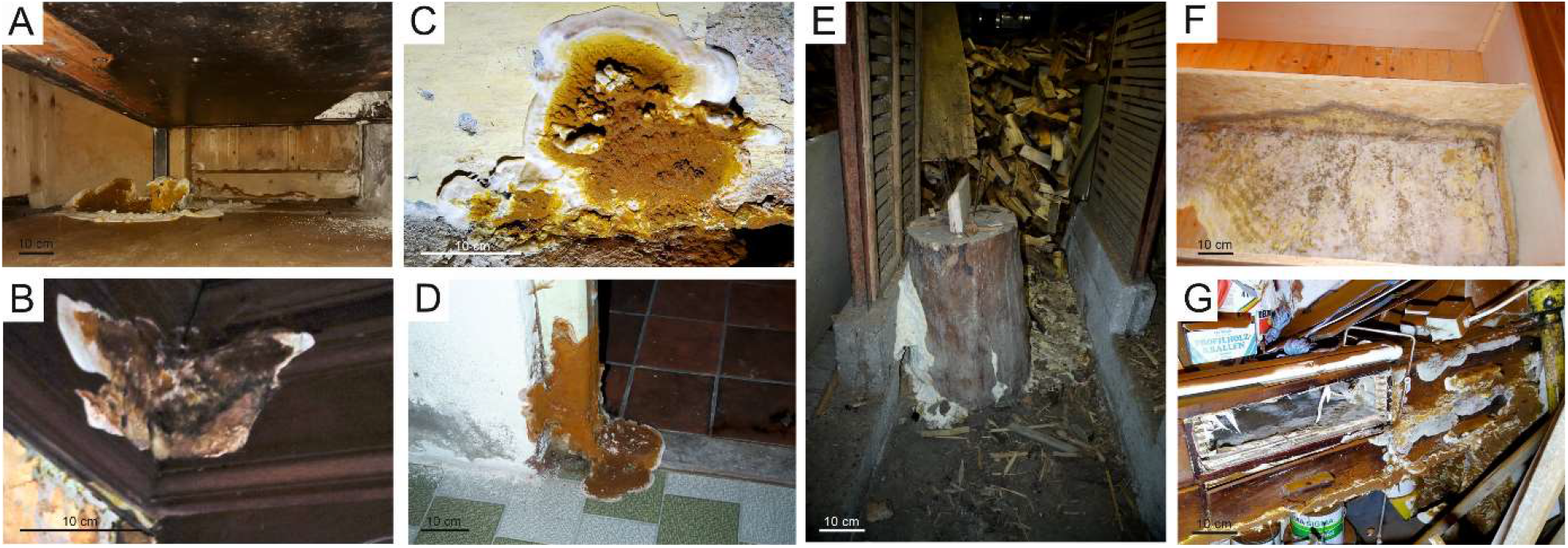
Fruiting bodies of *S. lacrymans* from distinct sampling points and examples for infestation of timber. (**A**) Zirl, (**B**) Reith/Alpbachtal, (**C**) Wolfsegg/Hausruck, (**D**) Grins, and (**E**) Tarrenz. (**F**) mycelium mat of *S. lacrymans*, (**F**) and (**G**) destruction caused by *S. lacrymans*.

The samples were collected and transferred into fresh zip-loc plastic bags. While wearing sterile gloves, the material was photo documented, small parts of the fruiting bodies and mycelium mats were cut out with a sterile scalpel, and transferred into 1.5 ml tubes filled with a sterile sphere matrix (0.2 g of glass beads, Ø 2 mm). Substrate material was transferred analogously into bead matrix tubes using a sterilized spoon, tweezers or a scalpel. The samples were stored at -20 °C until further processing.

### DNA Extraction and Sequencing

Total genomic DNA was extracted from the sampled material using the Dneasy^®^ PowerSoil^®^ Pro Kit (Qiagen, The Netherlands) according to the manufacturer’s instructions with noted alterations. Briefly, samples stored in the tubes containing the sphere matrix (1.5-ml tubes with 0.2 g of glass beads with 2 mm diameter) were frozen with liquid nitrogen and homogenized twice using FastPrep-24 5G (MP Biomedicals, Santa Anna, USA) at 6 m s^-1^ for 30 s (2 cycles, 300 sec pause).

Cells were treated with 800 µl CD1 buffer mixed with 10 µl Proteinase K solution (10 µg ml^-1^) and heated at 60 °C for 10 min after mixing. The tubes were horizontally secured on a Vortex adapter and vortexed for 10 min at maximum speed. Further steps were executed according to the kit’s instructions. In the last step, DNA was eluted from the columns using 50 µl of solution C6 (10 mM Tris). Three replicate extractions per sample were performed and the resulting DNA from all three extractions was pooled to represent the whole variability in our molecular fingerprint.

Sequencing was outsourced to Novogene Co., Ltd. (Beijing, China). The 16S rRNA gene variable regions V3–V4 were amplified using bacterial primers 341F (5′-CCTAYGGGRBGCASCAG-3′) and 806R (5′-GGACTACNNGGGTATCTAAT-3′). The samples were sequenced using the Illumina NovaSeq 6000 platform in rapid-mode paired end 250 bp (PE250). The sequences obtained from each sample were submitted to Sequence Read Archive and are available under BioProject PRJNA878418 (accession numbers SAMN30726529 to SAMN30726547).

### Statistics and analysis of sequencing data

Novogene Co., Ltd. (Beijing, China) was commissioned for sequencing data quality control and further metagenomic and statistical analysis, including α- and β-diversity. Our data was hence obtained using Novogene standard procedures that are briefly described below (for details see supplement).

#### a) Sequencing data processing

Paired-end reads were assigned to samples based on their unique barcodes and truncated by cutting off the barcode and primer sequences. They were merged using <u>FLASH</u> (V1.2.7) (Magoč and Salzberg 2011) (merged raw reads 30 k). Quality filtering of the raw tags followed the <u>Qiime</u> (V1.7.0) (Caporaso, Kuczynski et al. 2010) quality control process. The tags were compared with the reference database <u>SILVA138</u> (Quast, Pruesse et al. 2013) using <u>UCHIME</u> (Edgar, Haas et al. 2011) for chimera detection and removal.

#### b) OTU clustering and Taxonomic annotation

Sequences analysis was performed by <u>Uparse</u> (v7.0.1090) (Edgar 2013). Sequences with ≥97% similarity were assigned to the same OTUs. Each representative sequence was compared against the SSUrRNA database of <u>SILVA138</u> (Wang, Garrity et al. 2007) for species annotation at each taxonomic rank (threshold: 0.8∼1) (kingdom, phylum, class, order, family, genus, species) using <u>Qiime</u> (version 1.7.0, method Mothur). To obtain the phylogenetic relationship of all OTUs of the representative sequences <u>Muscle</u> (version 3.8.31) (Edgar 2004) was used. OTU abundance was normalized against the sample with the least sequences. Normalized data was used for subsequent analysis of alpha- and beta diversity.

### c) Alpha Diversity

Alpha diversity was assessed through community richness indices (<u>Chao1</u> and <u>ACE</u>) and community diversity indices (<u>Shannon</u> and <u>Simpson</u>). Sequencing depth was evaluated using (<u>the Good’s coverage)</u> and an index of phylogenetic diversity (<u>PD_whole_tree</u>). All indices were calculated with QIIME (Version 1.7.0) and visualised with R software [version 2.15.3 by R core Team (2018)].

### d) Beta Diversity

Beta diversity of weighted and unweighted unifrac-distances was calculated by QIIME software (version 1.7.0). Cluster analysis was preceded by principal component analysis (PCA). FactoMineR (Lê, Josse et al. 2008) and ggplot2 (Hadley 2016) package in R software (version 2.15.3) were used to reduce the dimension of the original variables. Principal Coordinate Analysis (PCoA) visualization was realized with WGCNA package (Langfelder and Horvath 2008), stat packages, and ggplot2 package in R software (version 2.15.3). Anosim, MRPP, and Adonis were performed by R software (Vegan package: anosim function, mrpp function and adonis function) (Jari, Gavin L. et al. 2013). AMOVA was calculated by mothur (Schloss, Westcott et al. 2009) using amova function. T_test and drawing were conducted by R software.

### Network analysis

Networks were constructed based on 16S rRNA gene sequence data. The molecular ecological network analysis (<u>MENA</u>) pipeline (Deng, Jiang et al. 2012) was used to analyse the networks. In brief, there were four main steps for network constructions: (i) original data collection (OTU table), (ii) data standardization (with centered log-ratio transformation), (iii) pairwise correlation/similarity estimation, and (iv) adjacent matrix formatting based on a random matrix theory method (Zhou, Deng et al. 2010). Default settings for data preparation and random matrix theory settings were used with noted alteration: the network was constructed using the OTUs present in at least 14 samples to increase the sensitivity of the analysis. To construct the network 0.76 was used as cutoff. ‘Greedy modularity optimization’ was the method for module separation. Indices to evaluate the features of the nodes were as follows: (i) degree - a node with higher degree means that it is highly connected with other nodes (that is, high degree = strong relationship with others); (ii) betweenness centrality (BC) = among-module connectivity (Pi; the parameter was used to indicate nodes connecting modules [connectors], Pi > 0.62); (iii) closeness centrality (CC) = within-module connectivity (Zi), referring to highly connected nodes within modules (module hubs), Zi > 2.5); (iv) important nodes to both the network and its own module coherence = network hub (Zi > 2.5, Pi > 0.62); and (v) peripherals—for nodes within the module but few outside connection, Zi < 2.5 and Pi < 0.62) (Olesen, Bascompte et al. 2007). The network data was obtained from the online platform MENA and was rearranged and analyzed with the open-source software <u>Cytoscape</u> (V3.0.X) (Shannon, Markiel et al. 2003). Network was filtered (degree between 5 and 81 inclusive) before cluster analysis. Clustering was obtained with plugin CytoCluster (V2.1.0) (Li, Li et al. 2017), an app for analysis and visualization of clusters from network. We used OH-PIN algorithm with default values for clustering. Clusters were manually evaluated to determine dominant taxa.

## Results

### Data structure

We sequenced 19 samples in total and obtained 91 000 to 118 000 reads per sample, of which 60% passed quality filtering. 98% of the filtered reads were above the Q20 threshold (Tab. S1). We found a minimum of 1244 OTUs and a maximum of 2193 OTUs per sample (average 1835 OTUs per sample); total reads per sample were on average 73262, of which approx. 15% were unique OTUs (Fig. S2). The rarefaction curves showed that sequencing depth was adequate for describing the microbial communities in our samples (Fig. S3A). The curves became flatter, even if we grouped the samples into origin classes (F = fruiting body, M = mycelium mats, WS = wooden substrate, and MS = mineral substrate) and hence just scarce species remain to be elusive (Fig. S3B).

### Bacterial abundance and community composition of fungal material and surrounding

OTUs were used to analyze differences in biodiversity and abundance of fungal fruiting bodies, mycelium mats, and substrate samples (wooden and mineral substrate). Bacteria were identified as the most diverse and abundant group of prokaryotes; archaea were found in low abundances and restricted diversity (Tab. S2 and Fig. S4). One mineral substrate and mycelium sample (Tarrenz) had with 39% and 49%, respectively, exceptionally high numbers of archaea OTUs (e.g. Halobacterota); in all other samples archaeal OTUs accounted for 0.09 to 7% of all OTUs. The most abundant bacterial phyla across all samples were Actinomycetota and Pseudomonadota (mean 44.63% and 33.36%), followed by Bacteroidota and Bacillota (7.9% and 4.66%), although again the sample from Tarrenz was a clear outlier (e.g. 30% of fruiting body or 27% of wood community was assigned to Bacteroidota, while 7.9% was the average). Other abundant phyla were Acidobacteriodota (G.FB1 with 14% and SM.WS with 19%) and Chloroflexota (WE.FB2 with 13% and WE.MS with 15%) (Tab. S2).

The most abundant bacterial phylum associated with *S. lacrymans* fruiting bodies was Pseudomonadota (38.4%), followed by Actinomycetota and Bacteroidota (35.7% and 5.9%). The mycelium mats were dominated by OTUs belonging to the Actinomycetota (47.5%), Halobacterota (25.3%), and Pseudomonadota (14.1%). Actinomycetota was the most abundant phylum in both environment samples (WS – 44.5% and MS – 45.1%), followed by Bacillota in wood (31.6%) and Pseudomonadota in mineral substrate samples (31.4%). With 30.4%, Pseudomonadota were abundant in wood. Other prominent phyla were Bacteroidota in wooden substrate (9.8%) and Halobacterota in mineral substrate material (9.9%) (Fig. 2A). On class level Gammaproteobacteria and Actinobacteria were found to dominate in fruiting bodies (33% and 30.8%), whereby Actinobacteria was most abundant on mycelium mats and the environmental samples (M - 44.5%, WS – 41.2%, and MS – 32.4%).

**Figure 2:**
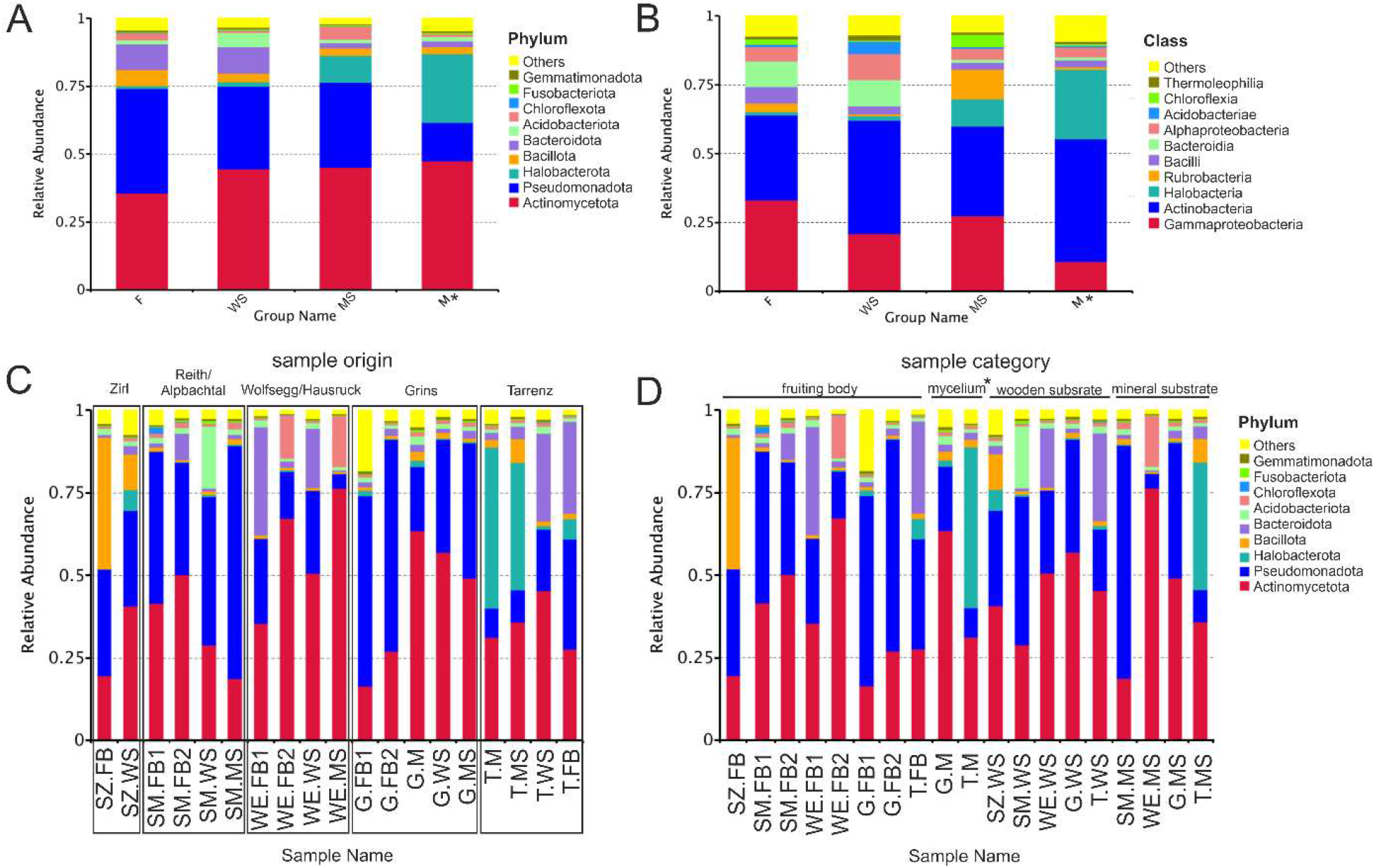
Prokaryote biodiversity and abundance is different in fungal fruiting bodies, mycelium mats, wooden substrate, and mineral substrate. The mean relative abundance (%) of ten most numerous bacterial (**A**) phyla, (**B**) classes. (**A+B**) based on sample type: F (fruiting body), WS (wooden substrate), MS (mineral substrate), and M (mycelium mats). (**C + D**) The mean relative abundance (%) of ten most numerous bacterial phyla in the individual samples (**C**) grouped based on sample origin and (**D**) grouped based on sample category (fruiting body, mycelium mats, wooden, and mineral substrate). * M (mycelium mats sample) n = 2, samples were included to display a trend. Graph is based on read numbers normalized by the total number of reads per each sample.

The prokaryotic community of woody and mineral substrates included Gammaproteobacteria (WS – 20.9% and MS – 27.4%). Gammaproteobacteria were also found in mycelium mats, but were represented by fewer reads (10.7%). Halobacteria were found associated with mycelium samples (25.3%) and mineral substrate material (9.9%). Another abundant class of prokaryotes in mineral substrate was Rubrobacteria (10.8%) (Fig. 2B).

Plotting of the 35 most abundant genera into a heatmap confirmed the trend visible in Fig. 2C and 2D, i.e. that there are distinct bacterial genera clusters for each sample type (Fig. 3A) and the clusters are about the same size (F – 11, M – 10, and WS - 10 members), except the cluster for mineral substrates, where just six members were detected. The bacterial community of the fruiting body was dominated by OTUs belonging to the phylum Pseudomonadota (55%), while Actinomycetota were dominant in samples derived from the mycelial mats (60%). Actinomycetota and Pseudomonadota emerged as abundant in mineral substrates and mycelial mats (33% and 40%). The genera *Advenella* sp. and *Pseudosphingobacterium* sp. were found to be similarly abundant in fruiting body and wood samples. Illustration of each individual sample shows clustering of samples from the same location, but also highlights pattern across sample type (Fig. 3B).

**Figure 3.**
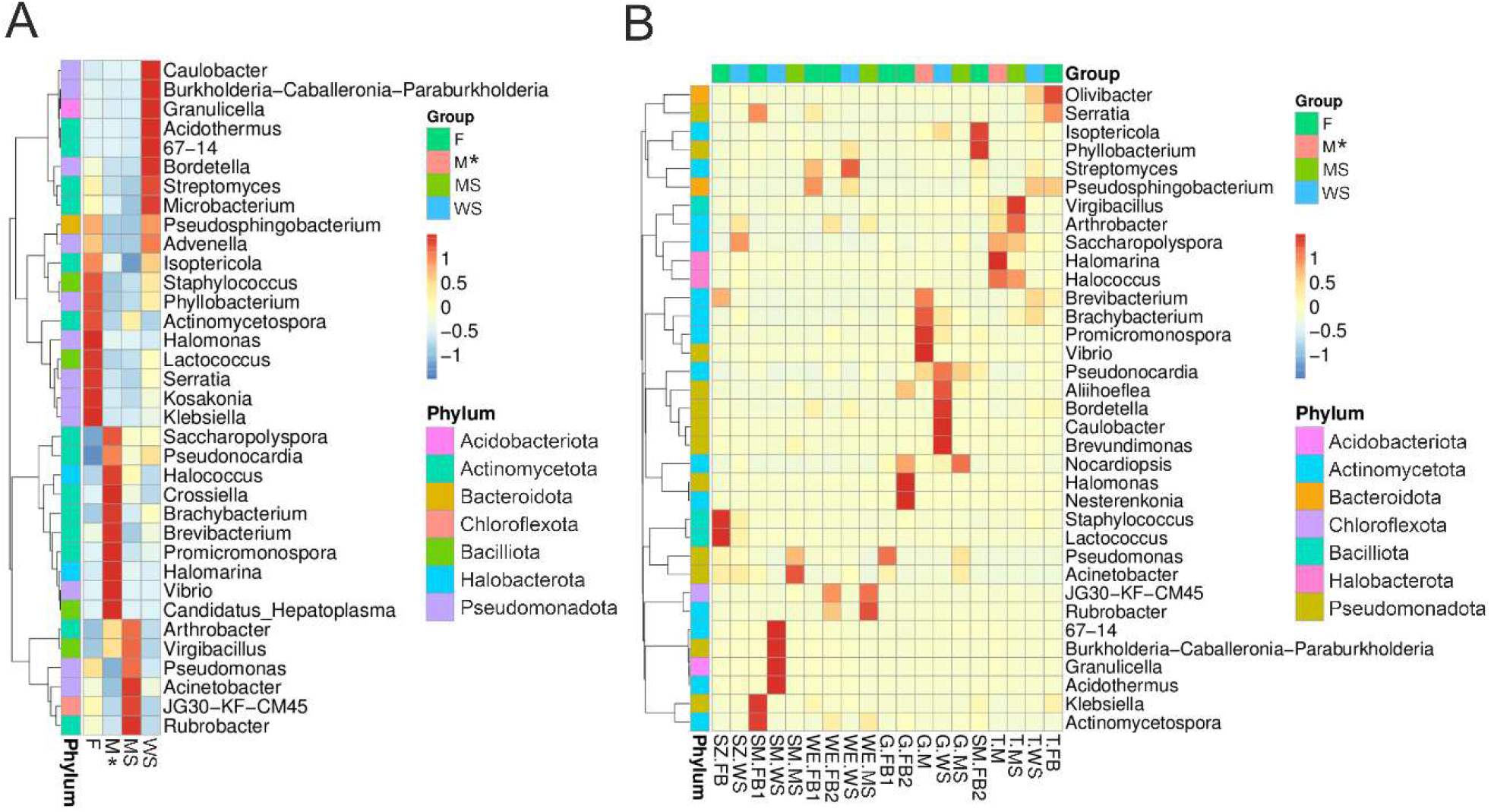
Taxonomic abundance cluster heat map (top 35 genera across samples) showing that prokaryote communities associated with different samples (mycelia, fruiting bodies, wood and mineral substrate) form distinct clusters. (**A**) Assignment of samples to groups (F= fruiting body, M = mycelium mats, MS = mineral substrate material, and WS = wood substrate), cumulative application. (**B**) Illustration of individual samples. The Y-axis represents the genus and the absolute value of ‘z’ (see color scheme red to blue) represents the distance between the raw score and the mean of the standard deviation. ‘Z’ is negative when the raw score is below the mean, and vice versa. Application by abundance. *M (mycelium mats sample) n = 2, samples were included for sake of completeness.

We used ternary plots to further screen for dominant taxa among the sample groups (fruiting body, wood, mineral substrate). Genera like *Pseudomonas, Acinetobacter*, and *Pseudonocardia* were evenly distributed over sample groups, while *Rubrobacter* and *Halococcus* were preferably found associated with the mineral substrate. Bacteria from the *Burkholderia-Caballeronia-Paraburkholderia* complex were identified on wood, while *Halomonas* sp. was found for fruiting body (Fig. S5 and Fig. S6). When the dataset of the ternary plots was subdivided according to sampling locations, genera patterns shift, as i.e for Grins, *Pseudonocardia* and *Pseudomonas* were more abundant compared to samples from Tarrenz or Wolfsegg/Hausruck (although they were not significantly assigned to one tissue/sample type) (Fig. S7A). Otherwise, the *Burkholderia-Caballeronia-Paraburkholderia* complex was found to be allocated to wood and *Acinetobacter* sp. and *Pseudomonas* sp. to mineral substrate from Reith/Alpbachtal (SM) (Fig. S7B). The material of the mineral substrate from Tarrenz was dominated by *Halococcus* sp.; another abundant genus was *Pseudosphingobacterium* sp. that was equally distributed between fruiting bodies and wood material (Fig. S7C). We found *Streptomyces* and *Pseudosphingobacterium* sp. as genera with major abundance in samples from Wolfsegg/Hausruck. *Rubrobacter* sp. was found associated to mineral substrate (Fig. S7D).

### Microbial Diversity

Alpha diversity indices including observed species, Chao1, Shannon index, phylogenetic diversity (PD) whole tree, and abundance-based coverage estimator (ACE) were calculated (Tab. S3). Chao-1 is a measure of total richness and is particularly useful because of a valid variance that is used to calculate confidence intervals (Chao 1987, Wang, Ahrné et al. 2005). The Shannon index reflects species numbers and evenness of species abundance (Guinane, Tadrous et al. 2013). Our results showed that the bacterial diversity indices were the highest for samples from mycelium mats (Fig. 4A-C), but these values should be interpreted with caution as only two distinct samples could be analyzed. Prokaryote communities associated with fruiting bodies were less diverse than those from other samples; however, these differences were not significant (Tab. S4). Overall, 1617 (30.6%) bacterial OTUs were found in all samples, while 732 OTUs (13%) were only present in samples from fruiting bodies, 451 OTUs in wood samples (8.5%) only, 374 OTUs solely in mineral substrates (7.1%), and 191 OTUs exclusively in the mycelial mats (3.6%) (Fig. 5A).

**Figure 4:**
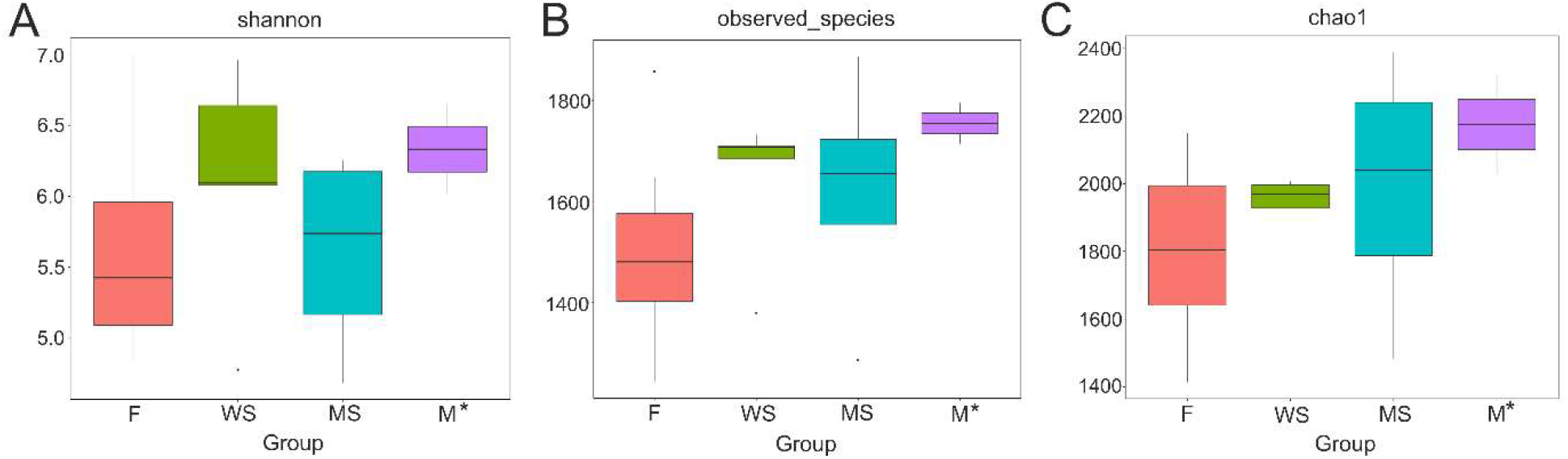
Alpha-diversity indices are more similar between mycelium mats and mineral-/ wood substrate samples than between samples from mycelium mats and fruiting body. Alpha diversity boxplots for differences in (**A**) Shannon Indexes, (**B**) observed species, and (**C**) Chao1 richness index. Fruiting body (F in salmon-colored), woody material (WS in green), mineral substrate samples (MS in blue), and fungal mycelium mats (M in turquoise). M (mycelium mats) * n = 2, samples were included for sake of completeness.

**Figure 5:**
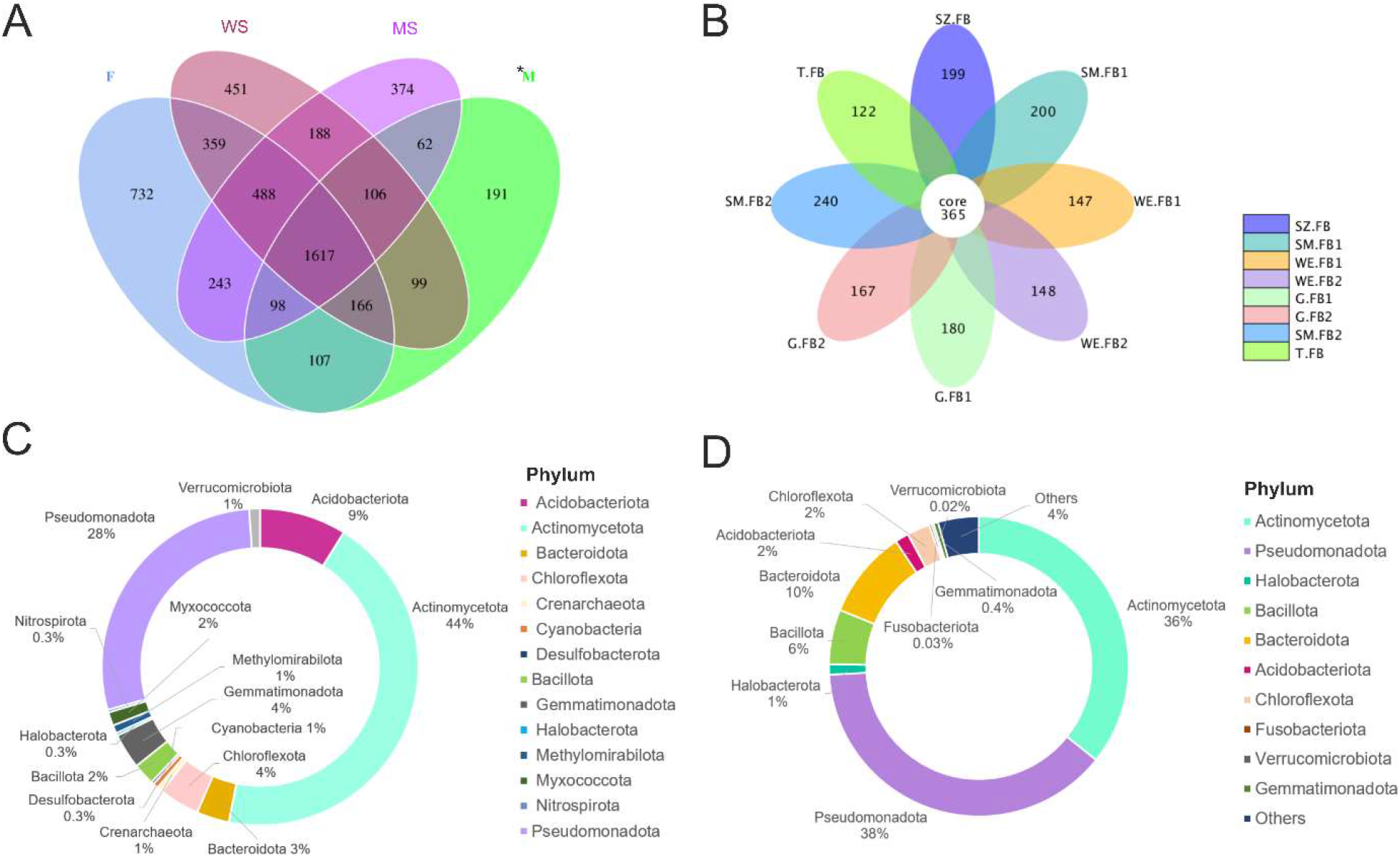
30% of OTU’s were shared across all samples and Actinomycetota dominated the core microbiome of *S. lacrymans* fruiting bodies. The numbers of unique and shared OTUs in the fruiting bodies (F), mycelium mats (M), and in the surrounding wood (WS) and mineral substrate (MS). (**A**) Venn diagram, (**B**) flower Diagram, (**C**) Core microbiome of *S. lacrymans* fruiting bodies, and (**D**) relative abundance of top 10 phyla associated with fruiting bodies of *S. lacrymans*. Different phyla are indicated with color code (legend right side). Note: M (mycelium sample) n = 2, samples were included for sake of completeness.

The core microbiome of *S. lacrymans* fruiting bodies analyzed in this study comprised 365 OTUs (Fig. 5B), while on average 175 unique OTUs were found per fruiting body. The fruiting body core microbiome was dominated by the genera *Nocardioides, Pseudomonas, Pseudonochardia, Streptomyces* and *Rubrobacter*. Rarer core OTUs belonged to *Solibacter, Saccharopolyspora, Vicinamibacteraceae, Sphingomonas*, and *Acinetobacter* (see supplement table 01_coreFB).

Core OTU numbers were dominated by Actinomycetota, as 160 (44%) of the 365 OTUs were assigned to this phylum. Another phylum with high OTU counts was Pseudomonadota (28%), followed by Acidobacteriota (9%), Chloroflexota, Gemmatimonadota (both 4%), Bacteriodota (3%), and Bacillota (2%) (Fig. 5C). Interestingly, core OTUs of fungal fruiting bodies included phyla from the domain Archaea, although these had very limited abundance (Crenarchaeota – 0.5% and Halobacterota - 0.3%). In comparison, the fruiting body pan microbiome was dominated by Pseudomonadota and Actinomycetota (38% and 36%), followed by Bacteriodota (10%) and Bacillota (6%) (Fig. 5D).

Beta diversity was analyzed with weighted-UniFrac analysis [Fig. S8AB (F – MS p = 0.95/ F – WS p= 0.64/ MS – WS p = 0.69 by Wilcox-test)]. Although, the differences in communities between distinct samples emerged as not significant, the β-diversity of the fruiting body sample from Grins (G.FB1) and mycelial and samples from the mineral substrate in Tarrenz (T.M and T.MS) was higher compared to other samples (Fig. S8A). Beta diversity of fruiting bodies was more similar to the one of wood samples than to that of samples from the mineral substrate. UniFrac-based PCoA provided an entire comparison of microbial communities and showed that the most distinct communities of wood material and samples from the mineral substrates were both more similar to communities of fruiting bodies than to each other. Communities of mycelial mats were more different to communities of other samples (Fig. S8C, caution n=2).

### Network analysis

An ecological network was constructed using the bacterial OTUs from the 19 dry rot - associated samples. This analysis resulted in a clustered network containing 415 nodes and 1141 edges (i.e. links) (Fig. 6A). The similarity threshold of the network was 0.76. Other indices, such as average connectivity, average path length, average clustering coefficient and modularity are summarized in Tab. S5. The network showed positive or negative interactions among taxa, whereby 184 of 1141 (16%) interactions were negative and 957 (84%) were positive (Tab. S5). A total of 6 clusters were identified, containing about 34% of the total OTUs: cluster 1 contained 65 OTUs, cluster 2 contained 30 OTUs, cluster 3 contained 18 OTUs, cluster 15 contained six OTUs, cluster 5 contained eleven OTUs, and cluster 6 contained three OTUs. Taxonomic classifications at phylum level in the network is shown in Fig. 6B. Generally, Actinomycetota, Pseudomonadota, Acidobacteriota were most abundant in the clusters, which is in accordance with their generally high abundance. Archaeal OTUs of the phyla Thermoproteota and Halobacterota were as well present in the network, although in minor amounts.

**Figure 6:**
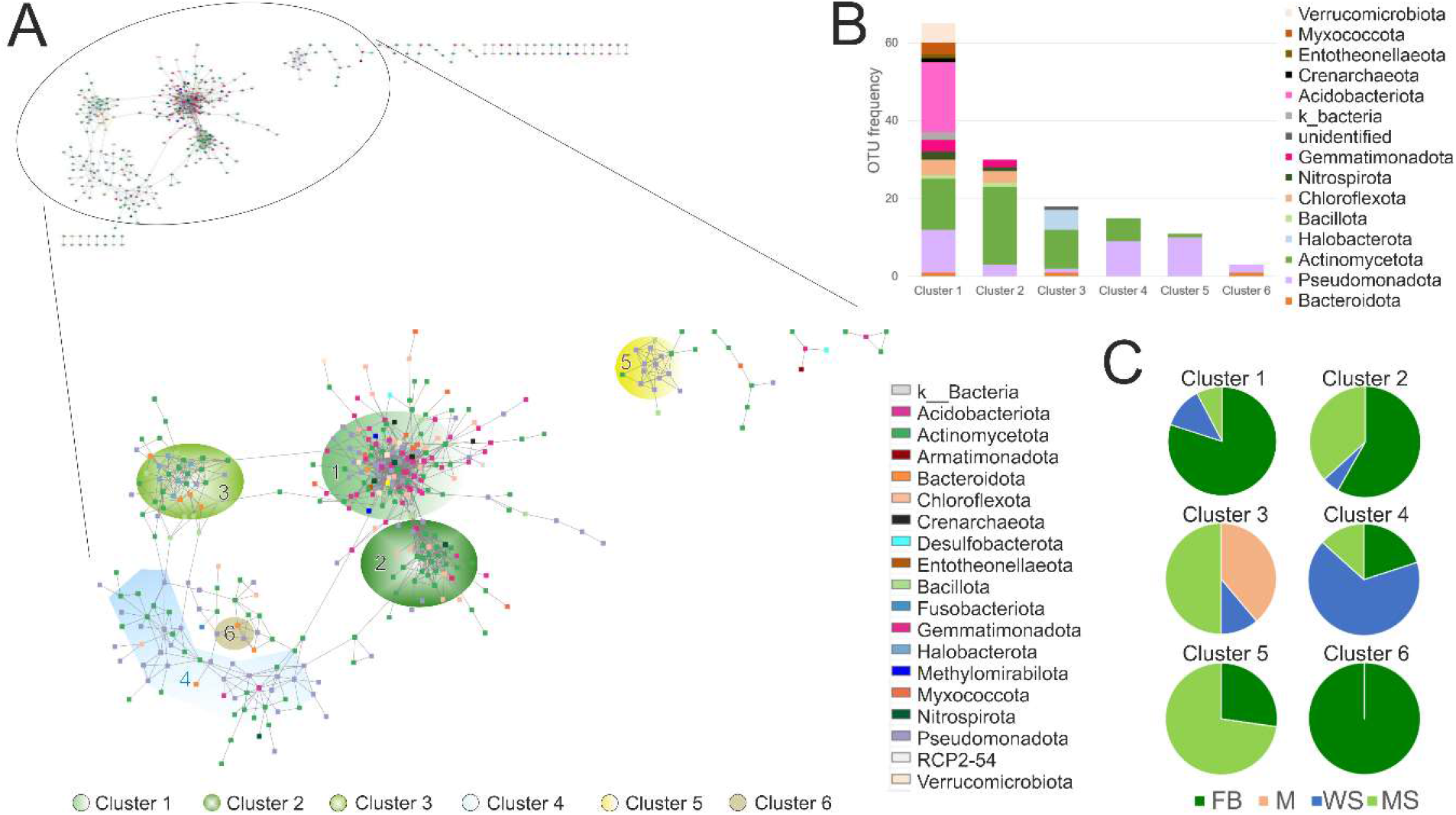
Microbial co-occurrence network showing clusters for distinct sample types. Each node (vertex) indicates a single OTU at 97% sequence similarity and different colors distinguish the nodes (i.e. OTUs) at the phylum level. (**A**) Network showing co-occurrence of OTUs, annotation to phyla (legend right bottom corner), and clustering. Left upper corner shows the entire network (415 nodes and 1141 edges), in the middle part an enlargement with clusters is displayed (149 nodes and 764 edges). (**B**) Bar plot showing OTU frequency (OTU richness) and taxonomic composition of clusters, and (**C**) Structure of clusters considering the main source of the OTUs. Fruiting bodies (F), mycelial mats (M), and in the surrounding wood (WS) and mineral substrates (MS). Note for mycelium n=2. For global network properties see Tab. S5.

Six OTUs were identified as cluster hubs, i.e. highly connected nodes within clusters with Zi > 2.5 (within-module connectivity (Zi) = closeness centrality), namely OTUs number 70 (Nitrospirota, *Nitrospira* sp.), 173 (Actinomycetota, order Gaiellales), 249 (Acidobacteriota, candidatus *Solibacter* sp.), 361 (Acidobacteriota, order Acidobacteriales), 390 (Actinomycetota, order Gaiellales), and 2822 (Pseudomonadota, family Xanthobacteraceae). All of them participate in cluster one, while OTU173 is as well found in cluster two. It was further analyzed if OTUs from clusters are predominantly occurring within a distinct sample (Fig. 6C). Clusters one, two, and six were predominantly comprised of OTUs mainly found associated with fruiting bodies (80%, 58%, and 100%), while cluster four was dominated by OTUs found mainly in wood (67%). Clusters three and five were predominantly comprised of OTUs mainly identified in underground material (50% and 73%, respectively). OTUs associated mainly with mycelium were exclusively found in cluster three.

## Discussion

Using an amplicon-based approach, the prokaryote communities associated with *S. lacrymans* were found to form tissue- and substrate-dependent clusters. The core community across all sample types was quite large with one third of the OTUs identified in all samples. The number of unique OTUs was small. Fruiting bodies of *S. lacrymans* shared a core set of 365 OTU’s, dominated by Actinomycetota, Pseudomonadota and Acidobacteriodota. Tissue/sample type emerged as the main driver for microbial diversity, followed by sample origin (Fig. 3). *Serpula lacrymans* forms resupinate fruiting bodies (which are adjacent to the substrate), hence it can be expected that the fruiting body community is clearly influenced by the community of the substrate on which the fruiting body has developed. Previous culture-based analyses of different *S. lacrymans* tissue types using a highly similar sampling approach have revealed marked differences: fruiting bodies and mycelia were dominated by Bacillota (formerly “Firmicutes”), rhizomorphs were dominated by Pseudomonadota (“Proteobacteria”), and in both tissues Actinomycetota and Bacteroidota (“Actinobacteria” and “Bacteroidetes”) were present but less abundant (Embacher, Neuhauser et al. 2021). Using amplicon sequencing, OTUs belonging to the same phyla were detected; however, compared to the culture-based studies their abundances were different. These differences are in parts the result of the complementary methodological approaches, as culture-based approaches favor different taxa compared to DNA-based approaches (Fonseca 2018, Embacher, Neuhauser et al. 2021). Also our amplicon approach did not address the issue of non-viable prokaryotes or exogenous DNA (Probst, Ascher-Jenull et al. 2021).

Nonetheless, the amplicon sequencing applied here is able to gain insight into abundance and structure of prokaryote communities. Combining the findings from amplicon-based diversity analyses and culture-based approaches allow to identify the most abundant bacterial phyla in fungal tissue and its surroundings and to get a first idea about the recruitment and specificity of selected taxa. Fruiting bodies showed the highest abundance (and diversity) of Pseudomonadota at the OTU level, while in culture-based studies Bacillota were the most abundant group (Embacher, Neuhauser et al. 2021). In mycelia, OTUs of Actinomycetota had a high abundance, while Bacillota dominated culture-based approaches. Culture-based experiments revealed that *Kluyvera* spp. (Pseudomonadota), *Paenisporosarcina* spp., *Bacillus* spp., and *Staphylococcus* spp. (all Bacillota) were most abundant on fruiting bodies of *S. lacrymans*. Another common genus was *Flavobacterium* spp. (Bacteriodetes). As some of the species which were abundant in culture dependent approaches form endospores (e.g. *Bacillus* spp.) it cannot be excluded that the high abundance of these species is biased by a stochastic ‘mixing’ and not a ‘true’ interaction. As OTU abundance and prevalence of taxa previously identified as dominating fruiting bodies was upstaged with amplicon sequencing, this hypothesis appears more likely (see supplement table 02_Top35taxa.xlsx). This reinforces that especially Bacillota, which were the most abundant endospore forming bacteria in isolation-based approaches, are likely not true interaction partners of *S. lacrymans*.

Distinct differences in the diversity and abundance of prokaryote taxa were found in the different sample types. OTU based clusters from fruiting bodies clearly separated from the mycelium, and vice versa. The biodiversity pattern from fungal tissue was more similar to the corresponding substrates than to each other, indicating that part of the *S. lacrymans* microbiome is recruited from the environment by stochastic processes (Fig. 3). An overlap in abundance of e.g. *Advenella* sp. and *Pseudosphingobacterium* sp. for fruiting body and wood samples was detected and some genera typically found associated with mineral substrates were as well present in fruiting bodies (e.g. *Pseudomonas* sp. or *Actinomycetospora* sp.). This indicates that bacteria are migrating from the substrate into fungal tissues and in developing fruiting bodies of *S. lacrymans* where only some selected species are able to persist. If this recruitment is the result of a random, stochastic event (Kielak, Scheublin et al. 2016) or if there is an environmental co-selection (Deveau, Bonito et al. 2018, Johnston, Hiscox et al. 2018) or if this is part of a specialized interaction (Boersma, Warmink et al. 2009, Christofides, Hiscox et al. 2019) remains to be investigated. The prokaryotic biodiversity found associated with the two samples from mycelial mats analyzed in this study was more dissimilar to the biodiversity associated with fruiting bodies than with the biodiversity of the corresponding substrates (Fig. 3). In contrast, alpha-diversity indices were more similar between mycelium mats and substrates than between mycelium mats and fruiting body. However, due to the small sample size for mycelial mats artefacts cannot be excluded, so more samples will be needed to draw robust conclusions.

The main link between OTUs in the clusters of the network was sample type, as e.g. 80% of OTUs found for cluster one are associated with fruiting bodies, and other clusters by OTUs associated with wood or mineral substrate (Fig. 6C). In other studies investigating fruiting body associated species, the relative abundance of bacterial groups varied between different parts of the fruiting bodies (e.g. pileus and stipe of *Morchella sextelata* or *Tricholoma matsutake)* and these communities differed from soil underneath the mushrooms (Oh, Kim et al. 2018, Benucci, Longley et al. 2019). As already mentioned, a comparison of these findings with those from resupinate fruiting bodies is limited. There is no differentiation between stipe and cap and the contact zone with the substrate embraces the lower surface of the entire fruiting body. Nonetheless, different fungal parts differ in tissue architecture and chemical composition, which could be responsible for the presence of different prokaryotic communities on the respective fungal tissue (Deja, Wieczorek et al. 2014). The main selectors for microbial diversity associated with *S. lacrymans* samples emerged to be sample type while the influence of the geographical origin of the samples seemed to be second to that. Similar findings have been reported for other fungi where climate and soil factors were negligible in comparison to host phylogeny (Gohar, Põldmaa et al. 2022). However, other studies argue that soil physical and chemical characteristics are important for the structure of mushroom-inhabiting bacterial communities at local scales (Pent, Põldmaa et al. 2017, Splivallo, Vahdatzadeh et al. 2019, Pent, Bahram et al. 2020). Based on all these findings, the here presented data suggests that also *S. lacrymans* has a core microbiome associated with different tissue types while an extensive accessory microbiome is the result of stochastic selection from the environment.

All samples analyzed here shared 30% of their OTUs. Alpha-diversity for fruiting bodies was lower than for the other samples. Interestingly, the number of unique total OTUs was highest in fruiting bodies. In previous culture-based studies, fruiting bodies had the lowest bacterial colony forming units per g (CFU g^-1^) tissue, but the highest phylogenetic diversity (Embacher, Neuhauser et al. 2021). We detected 365 OTUs as core microbiome of all *S. lacrymans* fruiting bodies. Those OTUs mainly belonged to the Actinomycetota and Pseudomonadota, followed by Acidobacteriodota. The identified *S. lacrymans* basidiomata core microbiome is very similar to the genera described to be the core global mushroom microbiome (Gohar, Põldmaa et al. 2022). The core mushroom microbiome comprises the genera *Halomonas, Serratia, Bacillus, Cutibacterium, Bradyrhizobium*, and *Burkholderia* (Gohar, Põldmaa et al. 2022), all of which, except *Bradyrhizobiuim*, were also detected in the core microbiome of fruiting bodies of *S. lacrymans*. Especially OTUs belonging to *Serratia* and *Halomonas* were abundant, while OTUs belonging to *Bacillus, Burkholderia*, and *Cutibacterium* were rare. Archaea OTUs (Crenarchaeota and Halobacterota) were also part of the fruiting body core microbiome, similar to findings in forest mushrooms (Rinta-Kanto, Pehkonen et al. 2018). The five most abundant phyla in the fruiting body core microbiome were Actinomycetota (44%), Pseudomonadota (28%), Acidobacteriota (9%), Chloroflexota (4%), and Gemmatimonadota (3.6%), while the most abundant phyla in the pan microbiome were Pseudomonadota and Actinomycetota (38% and 36%), followed by Bacteriodota (10%) and Bacillota (6%). In comparison to the core, the abundance of Acidobacteriota, Gemmatimonadota, and Chloroflexota diminished in the pan microbiome, while Bacillota and Bacteriodota were enriched, which could hint for a shift of secondary microbiota to primary microbiota.

The here revealed fruiting body core microbiome also contained cellulolytic and xylanolytic *Pseudonocardia* (Malfait, Godden et al. 1984, Kaewkla and Franco 2021), *Saccharopolyspora* spp. that produces xylanase (Sinma, Khucharoenphaisan et al. 2011), glycosyl transferases (Schmidt-Bohli 2016), cellulase (Meena, Rajan et al. 2013), and amylases (Chakraborty, Khopade et al. 2011, Xu, Liao et al. 2016), or *Sphingomonas* spp, which were shown to have an increased potential to use complex carbohydrates including cellulose, chitin, and hemicellulose (Tláskal and Baldrian 2021). Other species were *Pseudomonas* spp. and *Acinetobacter* spp., taxa which are known for their nitrogen fixing abilities (Liba, Ferrara et al. 2006, Wang, Bian et al. 2020, Yaghoubi Khanghahi, Strafella et al. 2021). Burkholderiales, a class for which a mycophagous lifestyle was suggested (Valášková, de Boer et al. 2009, Hervé, Ketter et al. 2016, Brabcová, Štursová et al. 2018, Starke, Morais et al. 2020), was as well quite abundant. Recent studies have identified distinct strategies of bacteria to fulfil nutritional aspects in deadwood and soil: (**i**) ‘decomposer’ species with high potential to degrade complex C-substrates (mainly bacteria from Acidobacteriota and Bacteroidota) and (**ii**) more ‘opportunistic’ bacteria that depend on labile or alternative carbon sources (mycophagy) (e.g. Pseudomonadota) (Lladó, Větrovský et al. 2019, Tláskal and Baldrian 2021). The latter were found to be more active in N-cycling, thereby providing a valuable task for the whole community (e.g. via N_2_ fixation) (Tláskal and Baldrian 2021). Nonetheless the physiological abilities of prokaryotes associated with *S. lacrymans* remain as open question and further research is needed to asses if and to what extent the prokaryotes detected in this study belong to the proposed groups of ‘opportunistic’ or ‘decomposer’ bacteria.

## Conclusion

This study provides further insights into the prokaryotic community associated with *S. lacrymans* and revealed that there is a core community associated with fungal fruiting bodies that is dominated by Actinomycetota and Pseudomonadota. Increasing information on deadwood bacteriomes and specific fungus-influencing microbiota hint at the importance of distinct microbes during wood decay, which potentially fulfil a precisely regulated set of tasks that help and influence others. In the case of timber decaying *S. lacrymans*, the exact succession and collaboration mechanisms (i.e. which taxa are helpful for wood decay of *S. lacrymans* and what is the underlying machinery*?*) are for the most part elusive. Here, we delivered further evidence of which species are associated with fruiting bodies of *S. lacrymans* and showed that the bacterial community of the ambient environment is distinct from that of the fungal tissue. As different fungal developmental stages or tissues have a distinct physiological set up and function, it is plausible that each phenotype could have specific microbial partners. Our results support this scenario as bacterial genera cluster are separated between fruiting bodies and mycelium. Additionally, assessment of the physiological properties of bacteria that are useful in the fungus/ wood environment (i.e. cellulolytic power, mycophagic behavior or involvement in nitrogen cycling) leave space for further research as e.g. bacteria from Actinomycetota are known for their ligninolytic abilities (Leo, Asem et al. 2018). Another recent publication assumed that phylogenetically different fungal hosts that fill the same ecological niche, harbor closely related bacterial taxa, and proposed that bacteria might prefer fungi with selected trophic modes (Robinson, House et al. 2021). Therefore, our study provides first puzzle pieces to resolve if fungi occupying and decaying indoor wood provide niches for closely related bacterial taxa, although further studies with fungi of similar trophic modes are necessary.

## Supporting information

supplement table 02_Top35taxa

supplement table 01_coreFB

## Acknowledgement

Doctoral program BioApp from Universität Innsbruck is acknowledged for funding. We also thank Universität Innsbruck for funding within the framework of the ‘Early Stage Funding’ (2021-Bio-1, no. 346314). SN was funded by the Austrian Science Fund: grant Y0801-B16.

## Conflict of interest

None declared.

## Notes

### Competing Interest Statement

The authors have declared no competing interest.

## References

Bahram, M., D. Vanderpool, M. Pent, M. Hiltunen and M. Ryberg (2018). “The genome and microbiome of a dikaryotic fungus (Inocybe terrigena, Inocybaceae) revealed by metagenomics.” Environmental Microbiology Reports 10(2): 155–166.

Benucci, G. M. N. and G. M. Bonito (2016). “The Truffle Microbiome: Species and Geography Effects on Bacteria Associated with Fruiting Bodies of Hypogeous Pezizales.” Microbial Ecology 72(1): 4–8.

Benucci, G. M. N., R. Longley, P. Zhang, Q. Zhao, G. Bonito and F. Yu (2019). “Microbial communities associated with the black morel Morchella sextelata cultivated in greenhouses.” PeerJ 7: e7744.

Berlemont, R. and A. C. Martiny (2013). “Phylogenetic Distribution of Potential Cellulases in Bacteria.” Applied and Environmental Microbiology 79(5): 1545–1554.

Boddy, L., J. C. Frankland and P. van West (2007). “Ecology of Saprotrophic Basidiomycetes.” Academic Press, London.

Boer, W., L. B. Folman, R. C. Summerbell and L. Boddy (2005). “Living in a fungal world: impact of fungi on soil bacterial niche development.” FEMS Microbiol Rev 29(4): 795–811.

Boersma, F. G. H., J. A. Warmink, F. A. Andreote and J. D. v. Elsas (2009). “Selection of Sphingomonadaceae at the Base of Laccaria proxima and Russula exalbicans Fruiting Bodies.” Applied and Environmental Microbiology 75(7): 1979–1989.

Boutelje, J. B. and A. F. Bravery (1968). Observations on the bacterial attack of piles supporting a Stockholm building. Stockholm, Svenska träforskningsinstitutet.

Brabcová, V., M. Štursová and P. Baldrian (2018). “Nutrient content affects the turnover of fungal biomass in forest topsoil and the composition of associated microbial communities.” Soil Biology and Biochemistry 118: 187–198.

Caporaso, J. G., J. Kuczynski, J. Stombaugh, K. Bittinger, F. D. Bushman, E. K. Costello, N. Fierer, A. G. Peña, J. K. Goodrich, J. I. Gordon, G. A. Huttley, S. T. Kelley, D. Knights, J. E. Koenig, R. E. Ley, C. A. Lozupone, D. McDonald, B. D. Muegge, M. Pirrung, J. Reeder, J. R. Sevinsky, P. J. Turnbaugh, W. A. Walters, J. Widmann, T. Yatsunenko, J. Zaneveld and R. Knight (2010). “QIIME allows analysis of high-throughput community sequencing data.” Nature Methods 7(5): 335–336.

Chakraborty, S., A. Khopade, R. Biao, W. Jian, X.-Y. Liu, K. Mahadik, B. Chopade, L. Zhang and C. Kokare (2011). “Characterization and stability studies on surfactant, detergent and oxidant stable α-amylase from marine haloalkaliphilic Saccharopolyspora sp. A9.” Journal of Molecular Catalysis B: Enzymatic 68(1): 52–58.

Chao, A. (1987). “Estimating the Population Size for Capture-Recapture Data with Unequal Catchability.” Biometrics 43(4): 783–791.

Christofides, S. R., J. Hiscox, M. Savoury, L. Boddy and A. J. Weightman (2019). “Fungal control of early-stage bacterial community development in decomposing wood.” Fungal Ecology 42: 100868.

Clausen, C. A. (1996). “Bacterial associations with decaying wood: a review.” International Biodeterioration & Biodegradation 37(1): 101–107.

Cowling, E. and W. Merrill (2011). “Nitrogen in wood and its role in wood deterioration.” Canadian Journal of Botany 44: 1539–1554.

Cullings, K., M. Stott, N. Marinkovich, J. DeSimone and S. Bhardwaj (2020). “Phylum-level diversity of the microbiome of the extremophilic basidiomycete fungus Pisolithus arhizus (Scop.) Rauschert: An island of biodiversity in a thermal soil desert.” MicrobiologyOpen.

Deja, S., P. P. Wieczorek, M. Halama, I. Jasicka-Misiak, P. Kafarski, A. Poliwoda and P. Młynarz (2014). “Do Differences in Chemical Composition of Stem and Cap of Amanita muscaria Fruiting Bodies Correlate with Topsoil Type?” PLOS ONE 9(12): e104084.

Deng, Y., Y.-H. Jiang, Y. Yang, Z. He, F. Luo and J. Zhou (2012). “Molecular ecological network analyses.” BMC Bioinformatics 13(1): 113.

Deveau, A., G. Bonito, J. Uehling, M. Paoletti, M. Becker, S. Bindschedler, S. Hacquard, V. Hervé, J. Labbé, O. A. Lastovetsky, S. Mieszkin, L. J. Millet, B. Vajna, P. Junier, P. Bonfante, B. P. Krom, S. Olsson, J. D. van Elsas and L. Y. Wick (2018). “Bacterial–fungal interactions: ecology, mechanisms and challenges.” FEMS Microbiology Reviews 42(3): 335–352.

Dutkiewicz, J., W. G. Sorenson, D. M. Lewis and S. A. Olenchock (1992). “Levels of bacteria, fungi and endotoxin in stored timber.” International Biodeterioration & Biodegradation 30(1): 29–46.

Edgar, R. C. (2004). “MUSCLE: multiple sequence alignment with high accuracy and high throughput.” Nucleic Acids Res 32(5): 1792–1797.

Edgar, R. C. (2013). “UPARSE: highly accurate OTU sequences from microbial amplicon reads.” Nature Methods 10(10): 996–998.

Edgar, R. C., B. J. Haas, J. C. Clemente, C. Quince and R. Knight (2011). “UCHIME improves sensitivity and speed of chimera detection.” Bioinformatics 27(16): 2194–2200.

Embacher, J., S. Neuhauser, S. Zeilinger and M. Kirchmair (2021). “Microbiota Associated with Different Developmental Stages of the Dry Rot Fungus Serpula lacrymans.” Journal of Fungi 7(5): 354.

Embacher, J., S. Zeilinger, M. Kirchmair, L. M. Rodrigues-R and S. Neuhauser (in press). “Wood decay fungi and their bacterial interaction partners in the built environment – a systematic review on fungal bacteria interactions in dead wood and timber.” Fungal Biology Reviews.

Folman, L., P. Gunnewiek, L. Boddy and W. de Boer (2008). “Impact of white-rot fungi on numbers and community composition of bacteria colonizing beech wood from forest soil.” FEMS microbiology ecology 63: 181–191.

Fonseca, V. G. (2018). ““Pitfalls in relative abundance estimation using eDNA metabarcoding”.” Molecular Ecology Resources 18(5): 923–926.

Gaylarde, C. C. and L. H. G. Morton (1999). “Deteriogenic biofilms on buildings and their control: A review.” Biofouling 14(1): 59–74.

Gohar, D., K. Põldmaa, L. Tedersoo, F. Aslani, B. Furneaux, T. W. Henkel, I. Saar, M. E. Smith and M. Bahram (2022). “Global diversity and distribution of mushroom-inhabiting bacteria.” Environmental Microbiology Reports n/a(/a).

Greaves, H. (1971). The bacterial factor in wood decay.

Guinane, C. M., A. Tadrous, F. Fouhy, C. A. Ryan, E. M. Dempsey, B. Murphy, E. Andrews, P. D. Cotter, C. Stanton, R. P. Ross and K. Bush (2013). “Microbial Composition of Human Appendices from Patients following Appendectomy.” mBio 4(1): e00366–00312.

Hadley, W. (2016). ggplot2: Elegant Graphics for Data Analysis Springer.

Hervé, V., E. Ketter, J.-C. Pierrat, E. Gelhaye and P. Frey-Klett (2016). “Impact of Phanerochaete chrysosporium on the Functional Diversity of Bacterial Communities Associated with Decaying Wood.” PLOS ONE 11(1): e0147100.

Hyde, K. D., A. M. S. Al-Hatmi, B. Andersen, T. Boekhout, W. Buzina, T. L. Dawson, D. C. Eastwood, E. B. G. Jones, S. de Hoog, Y. Kang, J. E. Longcore, E. H. C. McKenzie, J. F. Meis, L. Pinson-Gadais, A. R. Rathnayaka, F. Richard-Forget, M. Stadler, B. Theelen, B. Thongbai and C. K. M. Tsui (2018). “The world’s ten most feared fungi.” Fungal Diversity 93(1): 161–194.

Jari, O., S. Gavin L., B. F. Guillaume, K. Roeland, L. Pierre, M. Peter R., O. H. R.B. S. Peter, S. M. Henry H., S. Eduard, W. Helene, B. Matt, B. Michael, B. Ben, B. Daniel, C. Gustavo, C. Michael, D. C. Miquel, D. Sebastien, E. Heloisa Beatriz Antoniazi, F. Rich, F. Michael, F. Brendan, H. Geoffrey, O. H. Mark, L. Leo, M. Dan, O. Marie-Helene, R. C. Eduardo, S. Tyler, S. Adrian, T. B. Cajo J.F. and W. James (2013). “Package ‘vegan’ Community ecology package.”

Johnston, S. R., L. Boddy and A. J. Weightman (2016). “Bacteria in decomposing wood and their interactions with wood-decay fungi.” FEMS Microbiology Ecology 92(11).

Johnston, S. R., J. Hiscox, M. Savoury, L. Boddy and A. J. Weightman (2018). “Highly competitive fungi manipulate bacterial communities in decomposing beech wood (Fagus sylvatica).” FEMS Microbiology Ecology 95(2).

Kaewkla, O. and C. M. M. Franco (2021). “Pseudonocardia pini sp. nov., an endophytic actinobacterium isolated from roots of the pine tree Callitris preissii.” Archives of Microbiology 203(6): 3407–3413.

Kielak, A. M., T. R. Scheublin, L. W. Mendes, J. A. van Veen and E. E. Kuramae (2016). “Bacterial Community Succession in Pine-Wood Decomposition.” Frontiers in Microbiology 7(231).

Kržišnik, D., B. Lesar, N. Thaler and M. Humar (2018). “Micro and material climate monitoring in wooden buildings in sub-Alpine environments.” Construction and Building Materials 166: 188–195.

Langfelder, P. and S. Horvath (2008). “WGCNA: an R package for weighted correlation network analysis.” BMC Bioinformatics 9(1): 559.

Lê, S., J. Josse and F. Husson (2008). “FactoMineR: An R Package for Multivariate Analysis.” Journal of Statistical Software 25(1): 1 – 18.

Leo, V. V., D. Asem Zothanpuia and B. P. Singh (2018). Chapter 13 - Actinobacteria: A Highly Potent Source for Holocellulose Degrading Enzymes. New and Future Developments in Microbial Biotechnology and Bioengineering. B. P. Singh, V. K. Gupta and A. K. Passari, Elsevier: 191–205.

Li, M., D. Li, Y. Tang, F. Wu and J. Wang (2017). “CytoCluster: A Cytoscape Plugin for Cluster Analysis and Visualization of Biological Networks.” International Journal of Molecular Sciences 18(9): 1880.

Li, Q., C. Chen, P. Penttinen, C. Xiong, L. Zheng and W. Huang (2016). “Microbial diversity associated with Tricholoma matsutake fruiting bodies.” Microbiology 85(5): 531–539.

Liba, C. M., F. I. S. Ferrara, G. P. Manfio, F. Fantinatti-Garboggini, R. C. Albuquerque, C. Pavan, P. L. Ramos, C. A. Moreira-Filho and H. R. Barbosa (2006). “Nitrogen-fixing chemo-organotrophic bacteria isolated from cyanobacteria-deprived lichens and their ability to solubilize phosphate and to release amino acids and phytohormones.” Journal of Applied Microbiology 101(5): 1076–1086.

Line, M. A. (2003). “A nitrogen-fixing consortia associated with the bacterial decay of a wooden pipeline.” Letters in Applied Microbiology 25: 220–224.

Liu, Y., Q. Sun, J. Li and B. Lian (2018). “Bacterial diversity among the fruit bodies of ectomycorrhizal and saprophytic fungi and their corresponding hyphosphere soils.” Scientific Reports 8(1): 11672.

Lladó, S., T. Větrovský and P. Baldrian (2019). “Tracking of the activity of individual bacteria in temperate forest soils shows guild-specific responses to seasonality.” Soil Biology and Biochemistry 135: 275–282.

Magoč, T. and S. L. Salzberg (2011). “FLASH: fast length adjustment of short reads to improve genome assemblies.” Bioinformatics 27(21): 2957–2963.

Malfait, M., B. Godden and M. J. Penninckx (1984). “Growth and cellulase production of Micromonospora chalcae and Pseudonocardia thermophila.” Annales de l’Institut Pasteur / Microbiologie 135(1, Supplement B): 79–89.

Meena, B., L. A. Rajan, N. V. Vinithkumar and R. Kirubagaran (2013). “Novel marine actinobacteria from emerald Andaman & Nicobar Islands: a prospective source for industrial and pharmaceutical byproducts.” BMC Microbiology 13(1): 145.

Normung, D. I. f. (2020). DIN 68800-4 Wood preservation - Part 4: Curative treatment of wood destroying fungi and insects and refurbishment DIN 68800-4:2020-12, Beuth: 32.

Oh, S.-Y., M. Kim, J. A. Eimes and Y. W. Lim (2018). “Effect of fruiting body bacteria on the growth of Tricholoma matsutake and its related molds.” PLOS ONE 13(2): e0190948.

Olesen, J. M., J. Bascompte, Y. L. Dupont and P. Jordano (2007). “The modularity of pollination networks.” Proceedings of the National Academy of Sciences 104(50): 19891–19896.

Pent, M., M. Bahram and K. Põldmaa (2020). “Fruitbody chemistry underlies the structure of endofungal bacterial communities across fungal guilds and phylogenetic groups.” The ISME Journal 14(8): 2131–2141.

Pent, M., K. Põldmaa and M. Bahram (2017). “Bacterial Communities in Boreal Forest Mushrooms Are Shaped Both by Soil Parameters and Host Identity.” Frontiers in microbiology 8: 836–836.

Probst, M., J. Ascher-Jenull, H. Insam and M. Gómez-Brandón (2021). “The Molecular Information About Deadwood Bacteriomes Partly Depends on the Targeted Environmental DNA.” Frontiers in Microbiology 12.

Purahong, W., T. Wubet, G. Lentendu, M. Schloter, M. J. Pecyna, D. Kapturska, M. Hofrichter, D. Krüger and F. Buscot (2016). “Life in leaf litter: novel insights into community dynamics of bacteria and fungi during litter decomposition.” Molecular Ecology 25(16): 4059–4074.

Quast, C., E. Pruesse, P. Yilmaz, J. Gerken, T. Schweer, P. Yarza, J. Peplies and F. O. Glöckner (2013). “The SILVA ribosomal RNA gene database project: improved data processing and web-based tools.” Nucleic Acids Res 41(Database issue): D590–596.

Rehbein, M., G. Koch, U. Schmitt and T. Huckfeldt (2013). “Topochemical and transmission electron microscopic studies of bacterial decay in pine (Pinus sylvestris L.) harbour foundation piles.” Micron 44: 150–158.

Rinta-Kanto, J. M., K. Pehkonen, H. Sinkko, M. V. Tamminen and S. Timonen (2018). “Archaea are prominent members of the prokaryotic communities colonizing common forest mushrooms.” Canadian Journal of Microbiology 64(10): 716–726.

Robinson, A. J., G. L. House, D. P. Morales, J. M. Kelliher, L. V. Gallegos-Graves, E. S. LeBrun, K. W. Davenport, F. Palmieri, A. Lohberger, D. Bregnard, A. Estoppey, M. Buffi, C. Paul, T. Junier, V. Hervé, G. Cailleau, S. Lupini, H. N. Nguyen, A. O. Zheng, L. J. Gimenes, S. Bindschedller, D. F. Rodrigues, J. H. Werner, J. D. Young, P. Junier and P. S. G. Chain (2021). “Widespread bacterial diversity within the bacteriome of fungi.” Communications Biology 4(1): 1168.

Rogers, G. M. and A. A. W. Baecker (1991). “Clostridium xylanolyticum sp. nov., an Anaerobic Xylanolytic Bacterium from Decayed Pinus patula Wood Chips.” International Journal of Systematic and Evolutionary Microbiology 41(1): 140–143.

Schloss, P. D., S. L. Westcott, T. Ryabin, J. R. Hall, M. Hartmann, E. B. Hollister, R. A. Lesniewski, B. B. Oakley, D. H. Parks, C. J. Robinson, J. W. Sahl, B. Stres, G. G. Thallinger, D. J. V. Horn and C. F. Weber (2009). “Introducing mothur: Open-Source, Platform-Independent, Community-Supported Software for Describing and Comparing Microbial Communities.” Applied and Environmental Microbiology 75(23): 7537–7541.

Schmidt-Bohli, I.-I. (2016). Molekularbiologische und biochemische Untersuchungen zu Saccharothrix espanaensis, Saccharopolyspora erythraea und Kitasatospora sp. 152608 Dissertation, Freiburg.

Schulz-Bohm, K., O. Tyc, W. de Boer, N. Peereboom, F. Debets, N. Zaagman, T. K. S. Janssens and P. Garbeva (2017). “Fungus-associated bacteriome in charge of their host behavior.” Fungal Genetics and Biology 102: 38–48.

Shannon, P., A. Markiel, O. Ozier, N. S. Baliga, J. T. Wang, D. Ramage, N. Amin, B. Schwikowski and T. Ideker (2003). “Cytoscape: a software environment for integrated models of biomolecular interaction networks.” Genome Res 13(11): 2498–2504.

Sinma, K., K. Khucharoenphaisan, V. Kitpreechavanich and S. Tokuyama (2011). “Purification and Characterization of a Thermostable Xylanase from Saccharopolyspora pathumthaniensis S582 Isolated from the Gut of a Termite.” Bioscience, Biotechnology, and Biochemistry 75(10): 1957–1963.

Splivallo, R., M. Vahdatzadeh, J. G. Maciá-Vicente, V. Molinier, M. Peter, S. Egli, S. Uroz, F. Paolocci and A. Deveau (2019). “Orchard Conditions and Fruiting Body Characteristics Drive the Microbiome of the Black Truffle Tuber aestivum.” Frontiers in Microbiology 10.

Standards, A. (2015). ÖNORM B 3802-4 Protection of timber used in buildings - Part 4: Control and remediation measures against fungal decay and insect attack.

Starke, R., D. Morais, T. Větrovský, R. López Mondéjar, P. Baldrian and V. Brabcová (2020). “Feeding on fungi: genomic and proteomic analysis of the enzymatic machinery of bacteria decomposing fungal biomass.” Environmental Microbiology 22(11): 4604–4619.

Tarkka, M. T., B. Drigo and A. Deveau (2018). “Mycorrhizal microbiomes.” Mycorrhiza 28(5-6): 403–409.

Tauber, J. P., R. Gallegos-Monterrosa, A. T. Kovacs, E. Shelest and D. Hoffmeister (2018). “Dissimilar pigment regulation in Serpula lacrymans and Paxillus involutus during inter-kingdom interactions.” Microbiology-Sgm 164(1): 65–77.

Team, R. C. (2018). R: A language and environment for statistical computing.. V. R Foundation for Statistical Computing, Austria.

Tláskal, V. and P. Baldrian (2021). “Deadwood-Inhabiting Bacteria Show Adaptations to Changing Carbon and Nitrogen Availability During Decomposition.” Frontiers in Microbiology 12.

Tláskal, V., V. Brabcová, T. Větrovský, M. Jomura, R. López-Mondéjar, L. M. Oliveira Monteiro, J. P. Saraiva, Z. R. Human, T. Cajthaml, U. Nunes da Rocha and P. Baldrian (2021). “Complementary Roles of Wood-Inhabiting Fungi and Bacteria Facilitate Deadwood Decomposition.” mSystems 6(1): e01078–01020.

Tornberg, K., E. Bååth and S. Olsson (2003). “Fungal growth and effects of different wood decomposing fungi on the indigenous bacterial community of polluted and unpolluted soils.” Biology and Fertility of Soils 37: 190–197.

Vahdatzadeh, M., A. Deveau and R. Splivallo (2015). “The Role of the Microbiome of Truffles in Aroma Formation: a Meta-Analysis Approach.” Applied and Environmental Microbiology 81(20): 6946.

Valášková, V., W. de Boer, P. J. A. Klein Gunnewiek, M. Pospíšek and P. Baldrian (2009). “Phylogenetic composition and properties of bacteria coexisting with the fungus Hypholoma fasciculare in decaying wood.” The Isme Journal 3: 1218.

Wang, M., S. Ahrné, B. Jeppsson and G. Molin (2005). “Comparison of bacterial diversity along the human intestinal tract by direct cloning and sequencing of 16S rRNA genes.” FEMS Microbiology Ecology 54(2): 219–231.

Wang, M., Z. Bian, J. Shi, Y. Wu, X. Yu, Y. Yang, H. Ni, H. Chen, X. Bian, T. Li, Y. Zhang, L. Jiang and Q. Tu (2020). “Effect of the nitrogen-fixing bacterium Pseudomonas protegens CHA0-ΔretS-nif on garlic growth under different field conditions.” Industrial Crops and Products 145: 111982.

Wang, Q., G. M. Garrity, J. M. Tiedje and J. R. Cole (2007). “Naive Bayesian classifier for rapid assignment of rRNA sequences into the new bacterial taxonomy.” Appl Environ Microbiol 73(16): 5261–5267.

Wójcik-Fatla, A., B. Mackiewicz, A. Sawczyn-Domańska, J. Sroka, J. Siwiec, M. Paściak, B. Szponar, K. Pawlik and J. Dutkiewicz (2022). “Timber-colonizing gram-negative bacteria as potential causative agents of respiratory diseases in woodworkers.” International Archives of Occupational and Environmental Health 95(6): 1179–1193.

Xu, Y., C.-H. Liao, L.-L. Yao, X. Ye and B.-C. Ye (2016). “GlnR and PhoP Directly Regulate the Transcription of Genes Encoding Starch-Degrading, Amylolytic Enzymes in Saccharopolyspora erythraea.” Applied and Environmental Microbiology 82(23): 6819–6830.

Yaghoubi Khanghahi, M., S. Strafella, I. Allegretta and C. Crecchio (2021). “Isolation of Bacteria with Potential Plant-Promoting Traits and Optimization of Their Growth Conditions.” Current Microbiology 78(2): 464–478.

Yurkov, A., D. Krüger, D. Begerow, N. Arnold and M. T. Tarkka (2012). “Basidiomycetous Yeasts from Boletales Fruiting Bodies and Their Interactions with the Mycoparasite Sepedonium chrysospermum and the Host Fungus Paxillus.” Microbial Ecology 63(2): 295–303.

Zhou, J., Y. Deng, F. Luo, Z. He, Q. Tu and X. Zhi (2010). “Functional Molecular Ecological Networks.” mBio 1(4): e00169–00110.

