## Supplementary Figures for "Prokaryote communities associated with different types of tissue formed and substrates inhabited by *Serpula lacrymans*"

Technikerstrasse 25

A-6020 Innsbruck

Mail:

### Methods (Novogene Co., Ltd. (Beijing, China)

#### A) Sequencing preparation

##### 1. Library Construction, Quality Control and Sequencing

PCR amplification of targeted regions was performed by using specific primers connecting with barcodes. The PCR products with proper size were selected by 2% agarose gel electrophoresis. Same amount of PCR products from each sample was pooled, end-repaired, A-tailed and further ligated with Illumina adapters. Libraries were sequenced on a paired-end Illumina platform to generate 250bp paired-end raw reads. The experimental procedures of DNA library preparation are shown as follows:

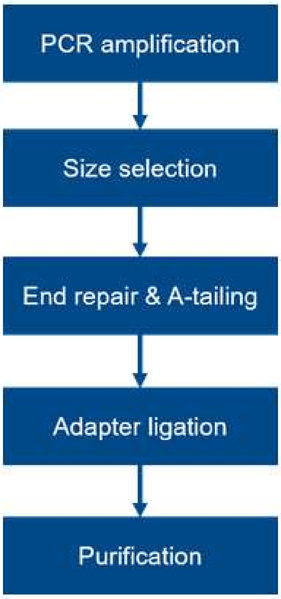

**Workflow of library construction**

The library was checked with Qubit and real-time PCR for quantification and bioanalyzer for size distribution detection. Quantified libraries will be pooled and sequenced on Illumina platforms, according to effective library concentration and data amount required.

#### B) Information analysis

##### 1 Sequencing data processing

Paired-end reads was assigned to samples based on their unique barcodes and truncated by cutting off the barcode and primer sequences. Paired-end reads were merged using FLASH (V1.2.7)[21] (see details http://ccb.jhu.edu/software/FLASH/), a very fast and accurate analysis tool, which was designed to merge paired-end reads when at least some of the reads overlap the read generated from the opposite end of the same DNA fragment, and the splicing sequences were called raw tags. Quality filtering on the raw tags were performed under specific filtering conditions to obtain the high-quality clean tags[22] according to the Qiime (V1.7.0)[23] (see details http://qiime.org/scripts/split_libraries_fastq.html) quality controlled process.
The tags were compared with the reference database (SILVA138 database, see details http://www.arb-silva.de/) using UCHIME algorithm (UCHIME Algorithm, see details http://www.drive5.com/usearch/manual/uchime_algo.html)[24] to detect chimera sequences (see details https://drive5.com/usearch/manual/chimeras.html). And then the chimera sequences were removed[25]. Then the Effective Tags finally obtained.

#### 2 OTU cluster and Taxonomic annotation

Sequences analysis were performed by Uparse software (Uparse v7.0.1090, see details http://drive5.com/uparse/)[26] using all the effective tags. Sequences with ≥97% similarity were assigned to the same OTUs. Representative sequence for each OTU was screened for further annotation.

For each representative sequence, Qiime (Version 1.7.0, see details http://qiime.org/scripts/assign_taxonomy.html)[27] in Mothur method was performed against the SSUrRNA database of SILVA138 Database (see details http://www.arb-silva.de/)[28]for species annotation at each taxonomic rank (Threshold:0.8~1)[29] (kingdom, phylum, class, order, family, genus, species).

To obtain the phylogenetic relationship of all OTUs representative sequences, the MUSCLE[30] (Version 3.8.31, see details http://www.drive5.com/muscle/) can compare multiple sequences rapidly.

OTUs abundance information were normalized using a standard of sequence number corresponding to the sample with the least sequences. Subsequent analysis of alpha diversity and beta diversity were all performed basing on this output normalized data.

#### 3 Alpha Diversity

Alpha diversity is applied in analysing complexity of biodiversity for a sample through 6 indices, including Observed-species, Chao1, Shannon, Simpson, ACE, Good-coverage. All these indices in our samples were calculated with QIIME (Version 1.7.0) and displayed with R software (Version 2.15.3).

Alpha Diversity Indices:

Community richness indices:

- Chao - the Chao1 estimator (see details http://scikit-bio.org/docs/latest/generated/skbio.diversity.alpha.chao1.html#skbio.diversity.alpha.chao1);
- ACE - the ACE estimator (see details http://scikit-bio.org/docs/latest/generated/skbio.diversity.alpha.ace.html#skbio.diversity.alpha.ace);

Community diversity indices:

- Shannon - the Shannon index (see details http://scikit-bio.org/docs/latest/generated/skbio.diversity.alpha.shannon.html#skbio.diversity.alpha.shannon);"
- Simpson - the Simpson index (see details http://scikit-bio.org/docs/latest/generated/skbio.diversity.alpha.simpson.html#skbio.diversity.alpha.simpson);

The index of sequencing depth:

- Coverage - the Good's coverage (see details http://scikit-bio.org/docs/latest/generated/skbio.diversity.alpha.goods_coverage.html#skbio.diversity.alpha.goods_coverage);

The index of phylogenetic diversity:

- PD_whole_tree - PD_whole_tree index (see details http://scikit-bio.org/docs/latest/generated/skbio.diversity.alpha.faith_pd.html?highlight=pd#skbio.diversity.alpha.faith_pd)

#### 4 Beta Diversity

Beta diversity analysis was used to evaluate differences of samples in species complexity, Beta diversity on both weighted and unweighted unifrac were calculated by QIIME software (Version 1.7.0). Cluster analysis was preceded by principal component analysis (PCA), which was applied to reduce the dimension of the original variables using the FactoMineR package and ggplot2 package in R software (Version 2.15.3). Principal Coordinate Analysis (PCoA) was performed to get principal coordinates and visualize from complex, multidimensional data. A distance matrix of weighted or unweighted unifrac among samples obtained before was transformed to a new set of orthogonal axes, by which the maximum variation factor is demonstrated by first principal coordinate, and the second maximum one by the second principal coordinate, and so on. PCoA analysis was displayed by WGCNA package, stat packages and ggplot2 package in R software (Version 2.15.3). Unweighted Pair-group Method with Arithmetic Means (UPGMA) Clustering was performed as a type of hierarchical clustering method to interpret the distance matrix using average linkage and was conducted by QIIME software (Version 1.7.0).

LEfSe analysis was conducted by LEfSe software. Metastat was calculated by R software. P-value was calculated by method of permutation test while q-value was calculated by method of Benjamini and Hochberg False Discovery Rate[31]. Anosim, MRPP and Adonis were performed by R software (Vegan package: anosim function, mrpp function and adonis function). AMOVA was calculated by mothur using amova function. T_test and drawing were conducted by R software.

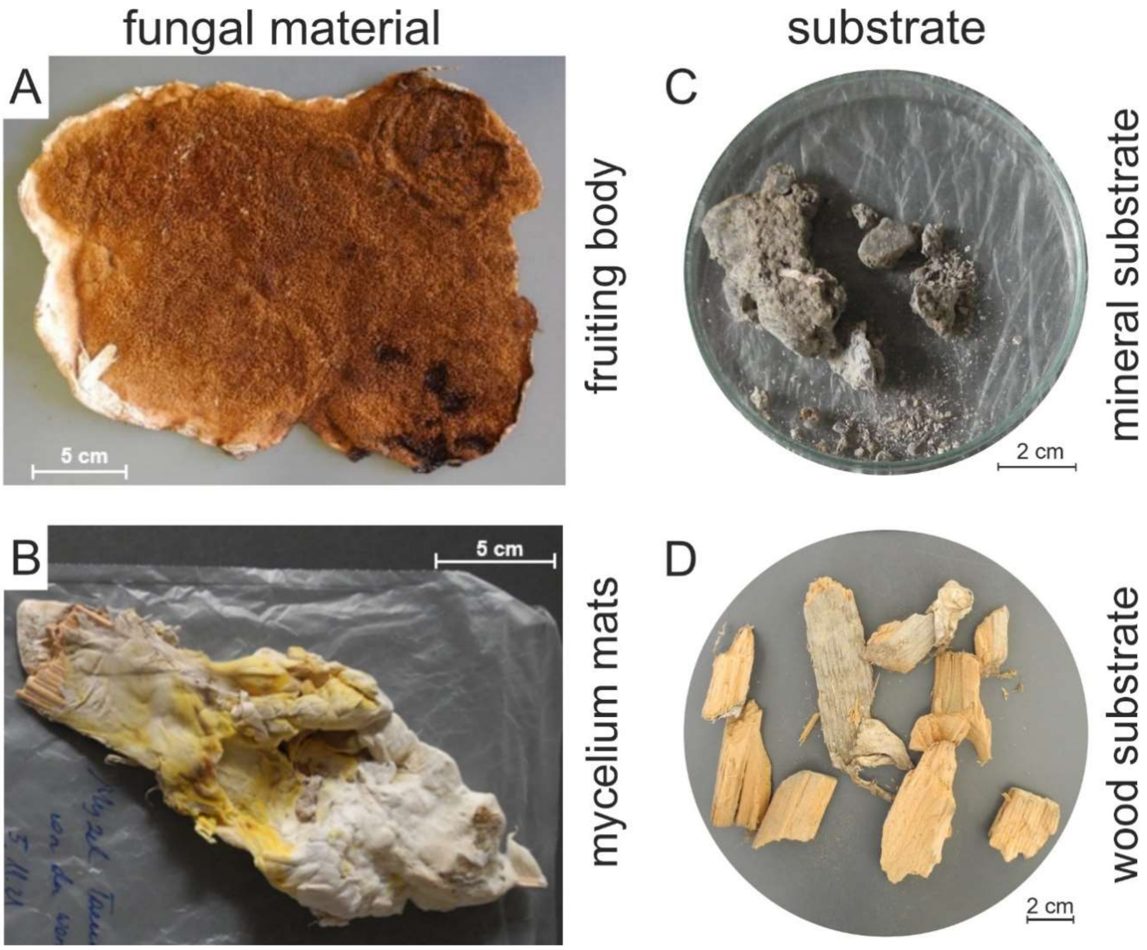

Figure S1: Samples used in this study. Fungal material (**A**) fruiting bodies and (**B**) mycelium mats of *S. lacrymans.* Substrate samples (**C**) mineral construction material (e.g. plaster, brick walls, or concrete) and (**D**) wood.

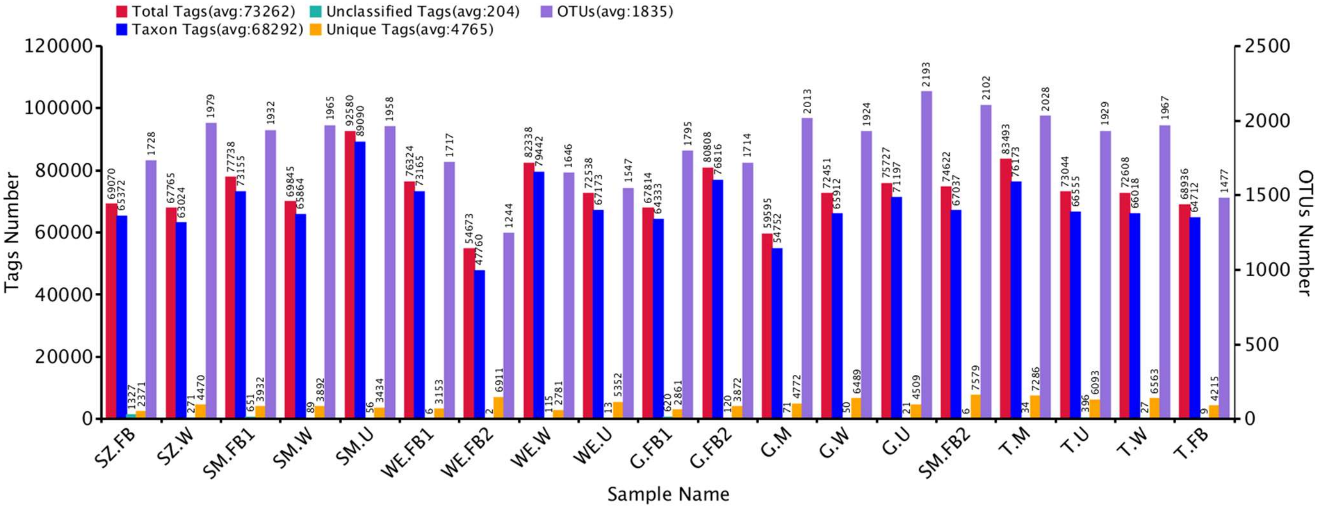

Figure S2: Summary of the tags and OTU numbers of each sample (SZ = Zirl, SM = Reith/Alpbachtal, WE = Wolfsegg am Hausruck, G = Grins, and T = Tarrenz / FB = Fruiting bodies, M = mycelium mats, W = wood, and U = MS mineral substrate sample).

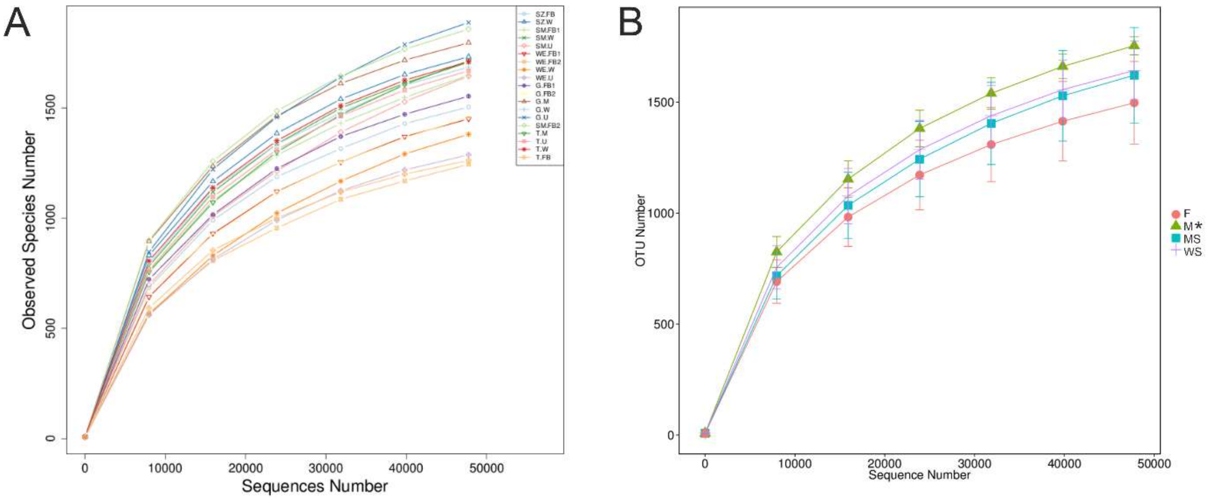

Figure S3: Rarefaction curves for fruiting body (FB, F), mycelium mats (M) (n=2), wood (WS), and mineral substrate (U = MS) samples. (**A**) Curves for all samples are separately shown (SZ = Zirl, SM = Reith/Alpbachtal, WE = Wolfsegg am Hausruck, G = Grins, and T = Tarrenz) and (**B**) curves for groups (in pale pink F = Fruiting bodies, in green M = mycelium, in violet WS = wood, and in blue MS mineral substrate). * M (mycelium mats) *n = 2, samples were included for sake of completeness.

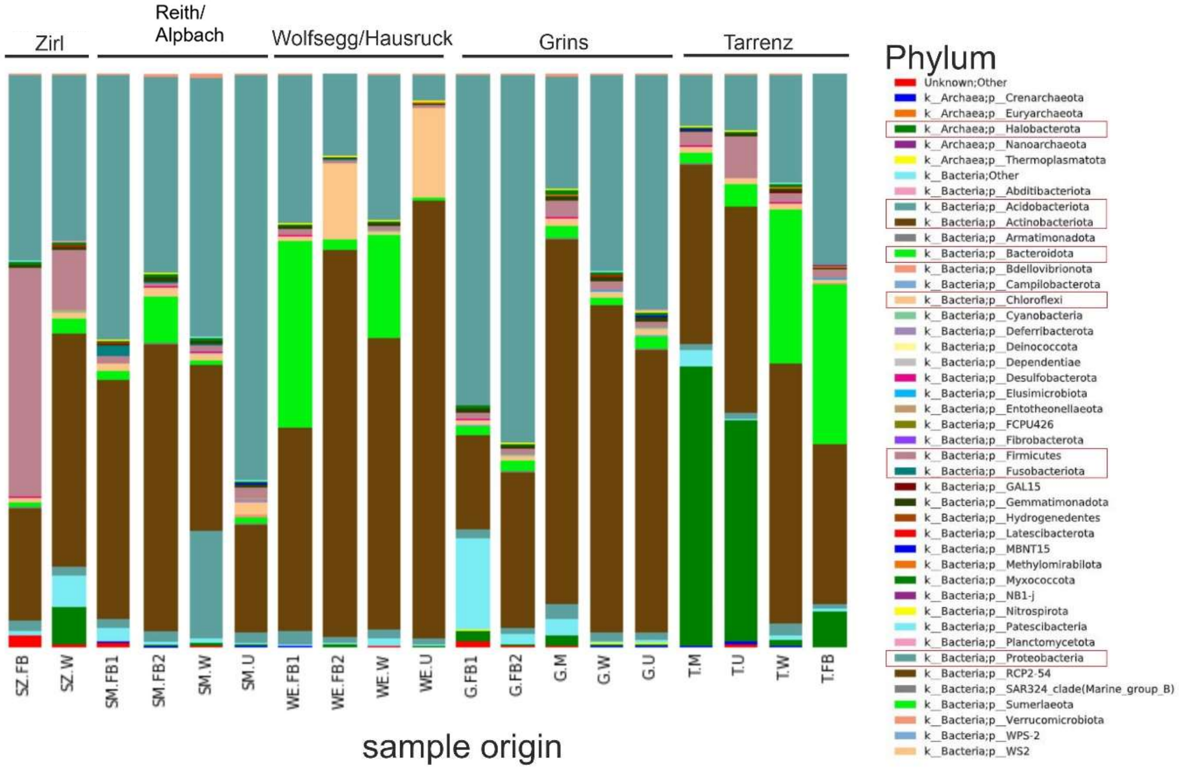

Figure S4: Taxa chart at phylum level. Grouping is based on sampling location (Zirl = SZ, Reith/Alpbachtal = SM, Wolfsegg/Hausruck = WE, Grins = G, Tarrenz = T). FB = fruiting body, M = mycelium mats, W = wood, and U = MS mineral substrate. Most abundant Phyla are marked with red boxes in the legend.

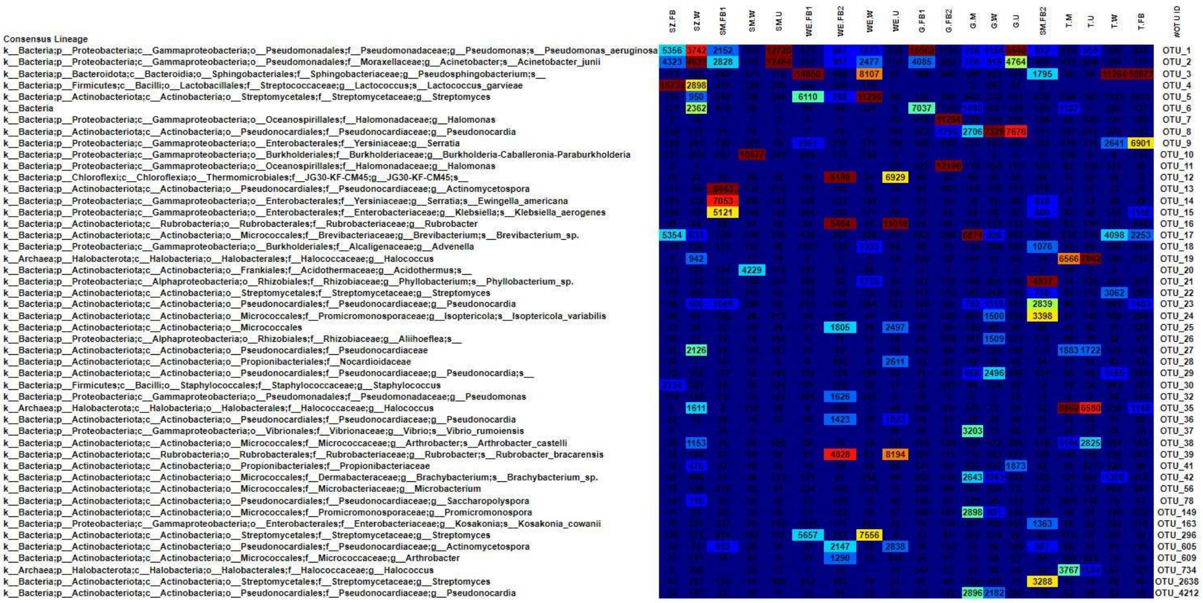

Figure S5: Heatmap of OTU sequence annotation results. OTUs with total counts of or above 3500 are displayed. The OTU heatmap displays raw OTU counts per sample; the counts are colored based on the contribution of each OTU to the total OTU count present in that sample (blue: contributes low percentage of OTUs to sample; red: contributes high percentage of OTUs).

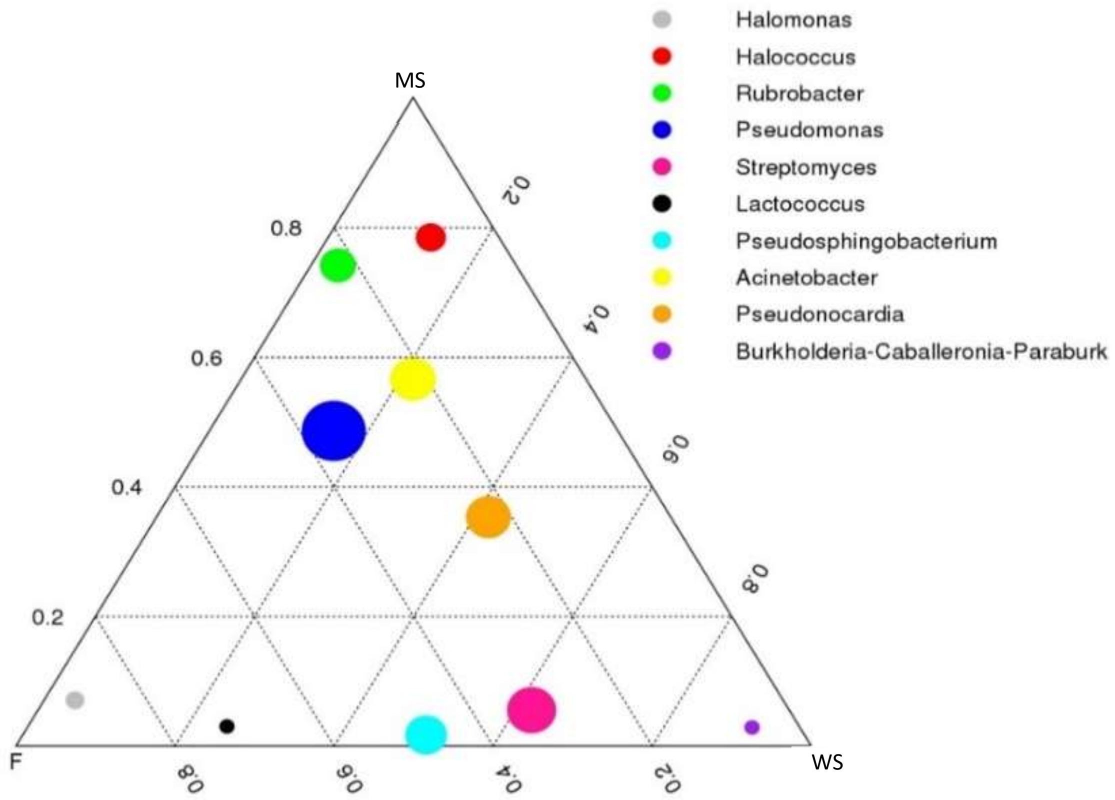

Figure S6: Ternary plot; the top 10 taxa with the average abundance of the three sample groups (Fruiting body – F, wood – WS, and mineral substrate sample – MS) at genus level were selected for generation. The three vertices in the graph represent three groups, the circles represent species, and the size of the circles is proportional to the relative abundance. The closer the circle is to a vertex, the higher the content of the species in the group.

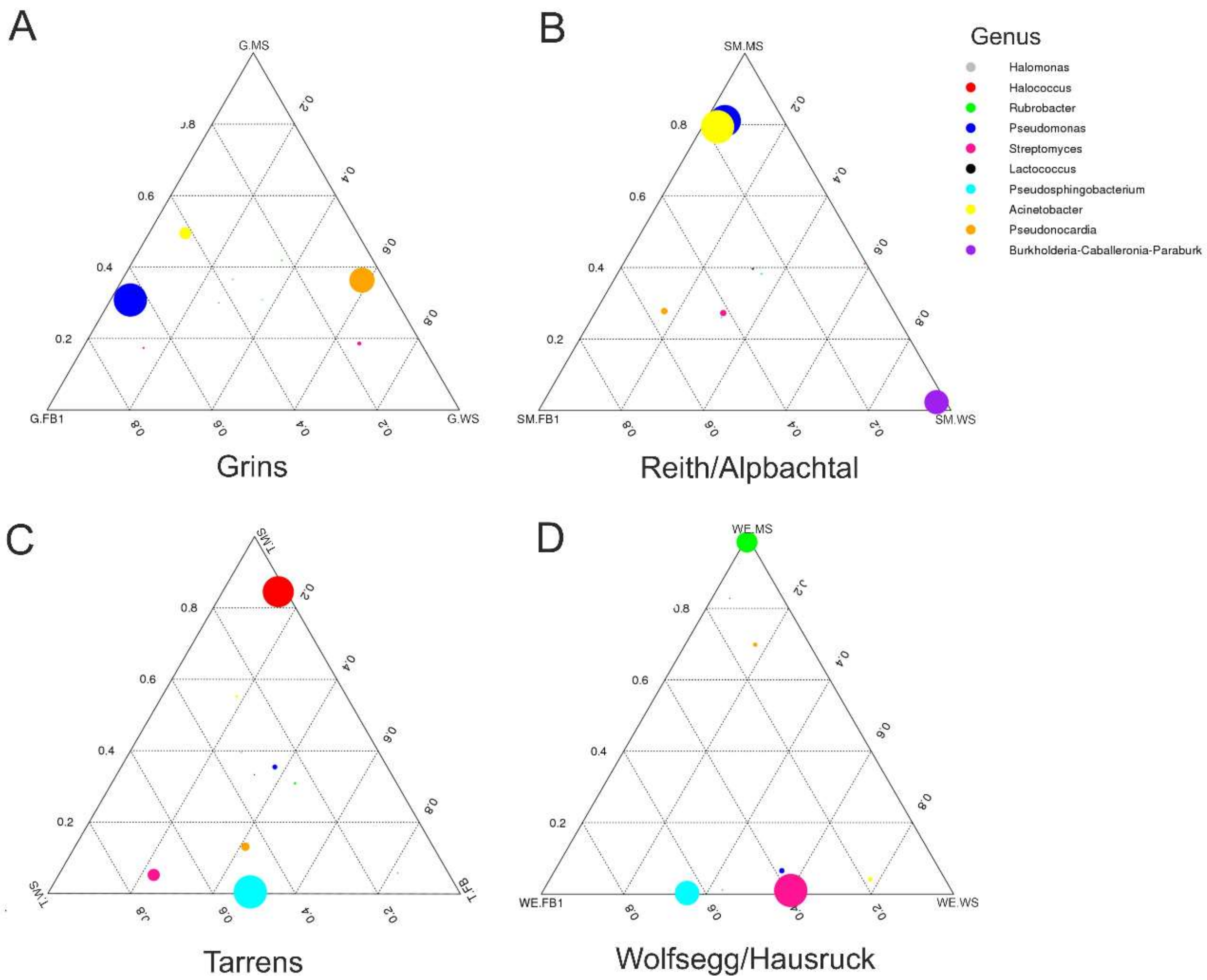

Figure S7: Genus ternary plots subdivided according to sampling locations (**A**) Grins, (**B**) Reith/Alpbachtal, (**C**) Tarrenz, and (**D**) Wolfsegg/Hausruck. Plot with the top 10 taxa with the average abundance of the three sample groups (Fruiting body – F, wood – WS, and mineral substrate sample – MS) at genus level were selected for generation. The three vertices in the graph represent three groups, the circles represent species, and the size of the circles is proportional to the relative abundance. The closer the circle is to a vertex, the higher the content of the species in the group. Samples from Zirl were excluded from this analysis, because underground sample was lacking.

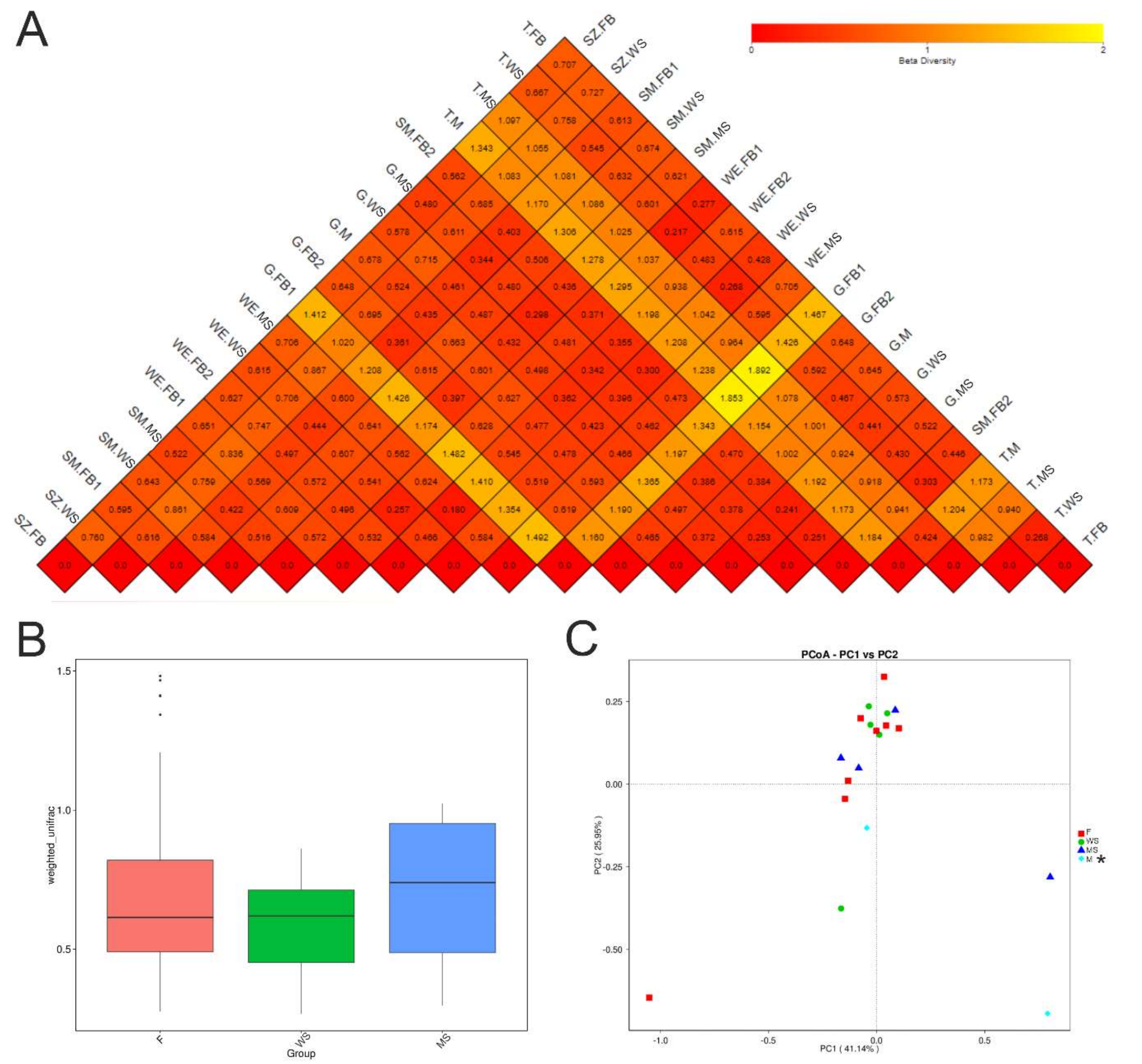

Figure S8: Analysis of beta diversity in bacterial communities of tissue from *S. lacrymans* compared with related wooden and mineral substrate. (**A**) Heatmap. Red color indicates no difference between samples; with increasing differences the brighter (orange or yellow) the boxes are. SZ = Zirl, SM = Reith/Alpbach, WE = Wolfsegg/Hausruck, G = Grins, T = Tarrenz; FB = fruiting body, M = mycelium, WS = wood, and MS = mineral substrate. (**B**) Histogram of beta diversity based on Unifrac distance matrices (F – MS p = 0.95/ F – WS p= 0.64/ MS – WS p = 0.69 by wilcox-*test*). (**C**) Unweighted principal coordinate analysis (PCoA – PC1 vs. PC2) representing the differences in bacterial community structure among fruiting body (F in red), woody material (WS in green), mineral substrate samples (MS in blue), and fungal mycelium (M in turquoise). M (mycelium mats) *n = 2, samples were included for sake of completeness.

**Table S1: Sequencing data processing summary** (SZ = Zirl, SM = Reith/Alpbachtal, WE = Wolfsegg am Hausruck, G = Grins, and T = Tarrenz / FB = Fruiting bodies, M = mycelium mats, WS = wooden, and MS = mineral substrate sample).

| Sample Name | Raw PE(#) | Raw Tags(#) | Clean Tags(#) | Effec-tive Tags(#) | Base(nt) | Avg Lene (nt) | Q20 | Q30 | GC% | Effec-tive % |
|---|---|---|---|---|---|---|---|---|---|---|
| G.FB1 | 112,216 | 86,451 | 84,357 | 67,814 | 28,626,323 | 422 | 98.29 | 94.37 | 52.83 | 60.43 |
| G.FB2 | 119,245 | 110,835 | 109,08 | 80,808 | 34,151,006 | 423 | 98.54 | 95.12 | 56.30 | 67.77 |
| G.M | 100,741 | 91,229 | 89,883 | 59,595 | 24,734,234 | 415 | 98.48 | 95.12 | 56.59 | 59.16 |
| G.MS | 105,018 | 101,63 | 100,247 | 75,727 | 31,739,802 | 419 | 98.42 | 94.86 | 56.03 | 72.11 |
| G.WS | 111,777 | 107,117 | 105,878 | 72,451 | 29,914,312 | 413 | 98.59 | 95.37 | 56.84 | 64.82 |
| SM. FB1 | 117,625 | 105,067 | 102,559 | 77,738 | 32,703,210 | 421 | 98.56 | 95.18 | 55.79 | 66.09 |
| SM. FB2 | 116,78 | 112,623 | 111,203 | 74,622 | 30,912,762 | 414 | 98.60 | 95.36 | 56.58 | 63.90 |
| SM.MS | 118,732 | 116,186 | 114,588 | 92,58 | 39,150,155 | 423 | 98.53 | 95.07 | 53.71 | 77.97 |
| SM.WS | 102,275 | 99,458 | 98,193 | 69,845 | 29,089,154 | 416 | 98.58 | 95.14 | 55.60 | 68.29 |
| SZ.FB | 102,296 | 93,74 | 90,348 | 69,07 | 29,204,054 | 423 | 98.25 | 94.34 | 52.95 | 67.52 |
| SZ.WS | 107,687 | 94,047 | 92,38 | 67,765 | 28,323,486 | 418 | 98.38 | 94.76 | 55.57 | 62.93 |
| T.FB | 105,846 | 99,755 | 98,525 | 68,936 | 28,890,681 | 419 | 98.56 | 95.25 | 55.20 | 65.13 |
| T.M | 118,694 | 111,183 | 109,55 | 83,493 | 33,741,184 | 404 | 98.33 | 94.61 | 59.00 | 70.34 |
| T.MS | 111,509 | 107,318 | 105,972 | 73,044 | 29,761,786 | 407 | 98.29 | 94.61 | 58.55 | 65.51 |
| T.WS | 111,133 | 106,918 | 105,467 | 72,608 | 30,279,419 | 417 | 98.53 | 95.21 | 55.61 | 65.33 |
| WE. FB1 | 110,962 | 108,239 | 106,69 | 76,324 | 31,947,708 | 419 | 98.53 | 95.19 | 54.83 | 68.78 |
| WE. FB2 | 91,041 | 88,144 | 87,041 | 54,673 | 22,815,425 | 417 | 98.54 | 95.27 | 57.98 | 60.05 |
| WE.MS | 103,005 | 100,306 | 99,039 | 72,538 | 30,362,150 | 419 | 98.51 | 95.18 | 58.55 | 70.42 |
| WE.WS | 109,344 | 102,486 | 101,128 | 82,338 | 34,232,960 | 416 | 98.62 | 95.34 | 55.46 | 75.30 |

**Notes:** Raw PE represents the original PE reads after sequencing; Raw Tags represents tags merged from PE reads; Clean Tags represents tags after filtering; Effective Tags represents tags after filtering chimera and can be finally used for subsequent analysis; Base is the number of bases of the Effective Tags; AvgLen represents average length of Effective Tags; Q20 and Q30 are the percentages of bases whose quality value in Effective Tags is greater than 20 (sequencing error rate is less than 1%) and 30 (sequencing error rate is less than 0.1%); GC (%) represents GC content in Effective Tags; Effective (%) represents the percentage of Effective Tags in Raw PE.

**Table S2: Community composition in percent [%] on phylum level.** Tissue types: Fruiting body (FB), mycelium mats (M), wood (WS), and mineral substrate (MS) samples. Sampling point: SZ = Zirl, SM = Reith/Alpbachtal, WE = Wolfsegg am Hausruck, G = Grins, and T = Tarrenz.

| | sample | | | | | | | | | | | | | | | | | | | |
|---|---|---|---|---|---|---|---|---|---|---|---|---|---|---|---|---|---|---|---|---|
| Phylum | Fruiting body | | | | | | | | wood substrate | | | | | mineral substrate | | | | mycelium | | **mean** |
| | SZ.FB | SM.FB1 | WE.FB1 | WE.FB2 | G.FB1 | G.FB2 | SM.FB2 | T.FB | SZ.WS | SM.Ws | WE.WS | G.WS | T.WS | SM.MS | WE.MS | G.MS | T.MS | G.M | T.M | |
| Bacillota | 41 | 1 | 1 | 0.7 | 1 | 1 | 0.6 | 2 | 11 | 1 | 0.8 | 2 | 2 | 2 | 0.5 | 1 | 12 | 3 | 5 | 4.66 |
| Pseudomonadota | 33 | 46 | 26 | 14 | 59 | 64 | 34 | 35 | 31 | 45 | 25 | 34 | 19 | 71 | 4 | 41 | 16 | 20 | 17 | 33.36 |
| Actinomycetota | 20 | 42 | 36 | 68 | 17 | 27 | 50 | 30 | 44 | 29 | 51 | 57 | 46 | 19 | 77 | 50 | 59 | 65 | 61 | 44.63 |
| Acidobacteriodota | 2 | 2 | 2 | 1 | 14 | 1 | 2 | 1 | 2 | 19 | 2 | 2 | 2 | 2 | 1 | 1 | 2 | 3 | 2 | 3.32 |
| Bacteriodota | - | 1 | 33 | 2 | 2 | 2 | 8 | 30 | 2 | 0.8 | 18 | 1 | 27 | 1 | 0.4 | 2 | 6 | 2 | 4 | 7.9 |
| Chloroflexota | - | - | - | 13 | - | - | 2 | 0.7 | 1 | 1 | 0.5 | 0.9 | 1 | 2 | 15 | 1 | 2 | 1 | - | 3.6 |
| Fusobacteriales | - | 2 | - | - | - | - | - | - | - | 0.02 | - | - | - | - | - | 0.01 | - | - | - | 0.68 |
| Archea | 0.09 | 0.08 | 0.2 | 0.5 | 2 | 0.3 | 0.4 | 6 | 7 | 0.7 | 0.2 | 0.6 | 1 | 0.6 | 0.3 | 0.7 | 39 | 2 | 49 | 5.82 |
| Species | | | | | | | | | | | | | | | | | | | | |
| *Microbacterium* spp. | 0.9 | 1 | 2 | 0.6 | 3 | 0.6 | 0.6 | 4 | 0.8 | 0.4 | 5 | 1 | 4 | 0.9 | 0.1 | 0.4 | 0.5 | 0.7 | 0.4 | 2.86 |

**Table S3: Alpha-Diversity Indices.** SZ = Zirl, SM = Reith/Alpbachtal, WE = Wolfsegg/Hausruck, G = Grins, T = Tarrenz; FB = fruiting body, M = mycelium mats, WS = woody substrate, and MS = mineral substrate.

| Sample name | observed_ species | shannon | simpson | chao1 | ACE | goods_ coverage | PD_whole_ tree |
|---|---|---|---|---|---|---|---|
| SZ.FB | 1504 | 4.840 | 0.855 | 1694.571 | 1817.958 | 0.992 | 187.781 |
| SZ.W | 1732 | 6.960 | 0.971 | 2006.542 | 2076.875 | 0.991 | 168.278 |
| SM.FB1 | 1648 | 5.909 | 0.927 | 2149.704 | 2127.455 | 0.989 | 255.671 |
| SM.WS | 1707 | 6.078 | 0.930 | 1996.093 | 2116.959 | 0.990 | 177.661 |
| SM.MS | 1644 | 5.327 | 0.859 | 2388.409 | 2432.866 | 0.986 | 138.804 |
| WE.FB1 | 1450 | 4.927 | 0.866 | 1824.513 | 1864.613 | 0.990 | 128.039 |
| WE.FB2 | 1244 | 6.115 | 0.955 | 1480.269 | 1635.753 | 0.992 | 114.320 |
| WE.WS | 1380 | 4.778 | 0.883 | 1928.008 | 1959.799 | 0.989 | 198.386 |
| WE.MS | 1287 | 4.685 | 0.885 | 1483.100 | 1611.116 | 0.992 | 119.426 |
| G.FB1 | 1553 | 5.147 | 0.848 | 1784.328 | 1890.523 | 0.991 | 196.116 |
| G.FB2 | 1458 | 5.180 | 0.873 | 1957.004 | 1996.352 | 0.990 | 155.859 |
| G.M | 1795 | 6.652 | 0.958 | 2027.081 | 2078.494 | 0.991 | 204.384 |
| G.WS | 1685 | 6.639 | 0.961 | 1927.286 | 2040.544 | 0.991 | 161.557 |
| G.MS | 1887 | 6.149 | 0.928 | 2190.680 | 2358.865 | 0.989 | 153.769 |
| SM.FB2 | 1857 | 7.000 | 0.969 | 2104.993 | 2208.114 | 0.990 | 174.372 |
| T.M | 1714 | 6.010 | 0.934 | 2323.122 | 2354.755 | 0.987 | 169.947 |
| T.MS | 1668 | 6.251 | 0.943 | 1889.187 | 2018.398 | 0.991 | 130.780 |
| T.WS | 1710 | 6.093 | 0.925 | 1969.216 | 2076.778 | 0.991 | 154.479 |
| T.FB | 1262 | 5.672 | 0.918 | 1413.048 | 1495.294 | 0.994 | 113.622 |

**Tab. S4: Wilcox Test output Alpha-Diversity.** a) Shannon, b) observed species, and c) chao1 richness index. FB = fruiting body, M = mycelium mats, W = wood, and MS = mineral substrate.

**a) Shannon**

| | Difference | pvalue | sig. | LCL | UCL |
|---|---|---|---|---|---|
| F - M | -5.125 | 0.2819 | | -14.911164 | 4.661164 |
| F - MS | -0.875 | 0.8090 | | -8.455330 | 6.705330 |
| F - W | -3.425 | 0.3173 | | -10.481903 | 3.631903 |
| M - MS | 4.250 | 0.4114 | | -6.470206 | 14.970206 |
| M - W | 1.700 | 0.7313 | | -8.656703 | 12.056703 |
| MS - W | -2.550 | 0.5227 | | -10.853836 | 5.753836 |

**b) observed species**

| | Difference | pvalue | sig. | LCL | UCL |
|---|---|---|---|---|---|
| F - M | -8.875 | 0.0497 | * | -17.734789 | -0.01521134 |
| F - MS | -3.375 | 0.3111 | | -10.237763 | 3.48776279 |
| F - W | -4.675 | 0.1397 | | -11.063884 | 1.71388446 |
| M - MS | 5.500 | 0.2458 | | -4.205412 | 15.20541221 |
| M - W | 4.200 | 0.3548 | | -5.176319 | 13.57631899 |
| MS - W | -1.300 | 0.7176 | | -8.817780 | 6.21777997 |

**c) chao1**

| | Difference | pvalue | sig. | LCL | UCL |
|---|---|---|---|---|---|
| F - M | -8.625 | 0.0602 | . | -17.668760 | 0.4187595 |
| F - MS | -4.125 | 0.2286 | | -11.130266 | 2.8802660 |
| F - W | -3.225 | 0.3086 | | -9.746548 | 3.2965477 |
| M - MS | 4.500 | 0.3483 | | -5.406942 | 14.4069422 |
| M - W | 5.400 | 0.2478 | | -4.171015 | 14.9710154 |
| MS - W | 0.900 | 0.8060 | | -6.773884 | 8.5738844 |

**Tab. S5: Global network properties** of Microbial co-occurrence network (Fig. 6).

| **Network Indexes** | **Microbial co-occurrence network (0.760)** |
|---|---|
| Total nodes | 415 |
| Total links | 1141 |
| - positive | 957 (≙ 84%) |
| - negative | 184 (≙ 16%) |
| R square of power-law | 0.913 |
| Average degree (avgK) | 5.499 |
| Average clustering coefficient (avgCC) | 0.251 |
| Average path distance (GD) | 5.853 |
| Geodesic efficiency (E) | 0.231 |
| Harmonic geodesic distance (HD) | 4.324 |
| Maximal degree | 56 |
| Nodes with max degree | OTU_390 |
| Centralization of degree (CD) | 0.123 |
| Maximal betweenness | 11453.944 |
| Nodes with max betweenness | OTU_127 |
| Centralization of betweenness (CB) | 0.127 |
| Maximal stress centrality | 447946 |
| Nodes with max stress centrality | OTU_643 |
| Centralization of stress centrality (CS) | 5.067 |
| Maximal eigenvector centrality | 0.288 |
| Nodes with max eigenvector centrality | OTU_390 |
| Centralization of eigenvector centrality (CE) | 0.269 |
| Density (D) | 0.013 |
| Reciprocity | 1 |
| Transitivity (Trans) | 0.297 |
| Connectedness (Con) | 0.582 |
| Efficiency | 0.981 |
| Hierarchy | 0 |
| Lubness | 1 |

### References

[1] Caporaso, J. Gregory, et al. Global patterns of 16S rRNA diversity at a depth of millions of sequences per sample. Proceedings of the National Academy of Sciences 108.Supplement 1 (2011): 4516-4522.

[2] Youssef, Noha, et al. Comparison of species richness estimates obtained using nearly complete fragments and simulated pyrosequencing-generated fragments in 16S rRNA gene-based environmental surveys. Applied and environmental microbiology 75.16 (2009): 5227-5236.

[3] Hess, Matthias, et al. Metagenomic discovery of biomass-degrading genes and genomes from cow rumen. Science 331.6016 (2011): 463-467.

[4] Asnicar F, Weingart G, Tickle T L, et al. Compact graphical representation of phylogenetic data and metadata with GraPhlAn[J]. PeerJ, 2015.

[5] DeSantis, T. Z., et al. NAST: a multiple sequence alignment server for comparative analysis of 16S rRNA genes. Nucleic acids research 34.suppl 2 (2006): W394-W399.

[6] Ondov, Brian D., Nicholas H. Bergman, and Adam M. Phillippy. Interactive metagenomic visualization in a Web browser. BMC bioinformatics 12.1 (2011): 385.

[7] Bulgarelli D, Garrido-Oter R, Münch P C, et al. Structure and function of the bacterial root microbiota in wild and domesticated barley[J]. Cell host & microbe, 2015, 17(3): 392-403.

[8] Li, Bing, et al. Characterization of tetracycline resistant bacterial community in saline activated sludge using batch stress incubation with high-throughput sequencing analysis. Water research 47.13 (2013): 4207-4216.

[9] Lundberg, Derek S., et al. Practical innovations for high-throughput amplicon sequencing.Nature methods 10.10 (2013): 999-1002.

[10] Lozupone, Catherine, and Rob Knight. UniFrac: a new phylogenetic method for comparing microbial communities. Applied and environmental microbiology 71.12 (2005): 8228-8235.

[11] Lozupone, Catherine, et al. UniFrac: an effective distance metric for microbial community comparison. The ISME journal 5.2 (2011): 169.

[12] Lozupone, Catherine A., et al. Quantitative and qualitative β diversity measures lead to different insights into factors that structure microbial communities. Applied and environmental microbiology 73.5 (2007): 1576-1585.

[13] Avershina, Ekaterina, Trine Frisli, and Knut Rudi. De novo Semi-alignment of 16S rRNA Gene Sequences for Deep Phylogenetic Characterization of Next Generation Sequencing Data. Microbes and Environments 28.2 (2013): 211-216.

[14] Magali Noval Rivas, PhD, Oliver T. Burton, et al. A microbita signature associated with experimental food allergy promotes allergic senitization and anaphylaxis. The Journal of Allergy and Clinical Immunology.Volume 131, Issue 1, Pages 201-212, January 2013.

[15] Anderson, M.J. 2001. A new method for non-parametric multivariate analysis of variance. Austral Ecology, 26: 32-46.

[16] McArdle, B.H. and M.J. Anderson. 2001. Fitting multivariate models to community data: A comment on distance-based redundancy analysis. Ecology, 82: 290-297.

[17] Warton, D.I., Wright, T.W., Wang, Y. 2012. Distance-based multivariate analyses confound location and dispersion effects. Methods in Ecology and Evolution, 3, 89-101.

[18] Zapala, M.A. and N.J. Schork. 2006. Multivariate regression analysis of distance matrices for testing associations between gene expression patterns and related variables. Proceedings of the National Academy of Sciences, USA, 103:19430-19435.

[19] Excoffier, L., Smouse, P.E. and Quattro, J.M. (1992) Analysis of molecular variance inferred from metric distances among DNA haplotypes: application to human mitochondrial DNA restriction data. Genetics, 131, 479-491.

[20] Segata, Nicola, et al. Metagenomic biomarker discovery and explanation. Genome Biol 12.6 (2011): R60.

[21] Magoč, Tanja, and Steven L. Salzberg. FLASH: fast length adjustment of short reads to improve genome assemblies. Bioinformatics 27.21 (2011): 2957-2963.

[22] Bokulich, Nicholas A., et al. Quality-filtering vastly improves diversity estimates from Illumina amplicon sequencing. Nature methods 10.1 (2013): 57-59.

[23] Caporaso, J. Gregory, et al. QIIME allows analysis of high-throughput community sequencing data. Nature methods 7.5 (2010): 335-336.

[24] Edgar, Robert C., et al. UCHIME improves sensitivity and speed of chimera detection. Bioinformatics 27.16 (2011): 2194-2200.

[25] Haas, Brian J., et al. Chimeric 16S rRNA sequence formation and detection in Sanger and 454-pyrosequenced PCR amplicons. Genome research 21.3 (2011): 494-504.

[26] Edgar, Robert C. UPARSE: highly accurate OTU sequences from microbial amplicon reads. Nature methods 10.10 (2013): 996-998.

[27] Altschul S F, Gish W, Miller W, et al. Basic local alignment search tool.[J]. Journal of Molecular Biology, 1990, 215(3):403-10.

[28] Wang, Qiong, et al. Naive Bayesian classifier for rapid assignment of rRNA sequences into the new bacterial taxonomy. Applied and environmental microbiology 73.16 (2007): 5261-5267.

[29] Quast C, Pruesse E, et al.The SILVA ribosomal RNA gene database project: improved data processing and web-based tools. Nucl. Acids Res. (2013) : D590-D596.

[30] MUSCLE: multiple sequence alignment with high accuracy and high throughputEdgar, 2004

[31] White, James Robert, Niranjan Nagarajan, and Mihai Pop. Statistical methods for detecting differentially abundant features in clinical metagenomic samples. PLoS computational biology 5.4 (2009): e1000352.
